# Approaches to Optimize Cell Internalization and *In Vivo* Tumor Homing by Aptamer-drug Conjugates using SELEX

**DOI:** 10.64898/2026.08.25.747018

**Authors:** Caroline D. Doherty, Sonia Jain, Katie K. Bakken, Brandon A. Wilbanks, Lauren L. Ott, Brett L. Carlson, Danielle M. Burgenske, Jann N. Sarkaria, L. James Maher

**Affiliations:** Department of Molecular Pharmacology and Experimental Therapeutics, Mayo Clinic Graduate School of Biomedical Sciences, Rochester, MN 55905; Department of Radiation Oncology, Mayo Clinic, Rochester, MN 55905; Department of Biochemistry and Molecular Biology, Mayo Clinic Graduate School of Biomedical Sciences, Rochester, MN 55905

**Keywords:** *in vivo* SELEX, conjugate SELEX, aptamer, aptamer drug conjugate, ApDC, MMAE, neuro-oncology, glioblastoma, GBM, PDX model, brain tumor, antibody drug conjugate, ADC

## Abstract

Glioblastoma (GBM) is the most common primary malignant brain tumor and is typically fatal. GBM therapies are hindered by the impermeability of the blood brain barrier (BBB), the diffuse and infiltrative nature of the tumor, and the high heterogeneity of intratumoral GBM cells. Aptamers are short, synthetic, folded single strands of RNA or DNA or analogs that bind targets with high affinity and specificity. Aptamers are developed via the principles of natural selection, permitting an unbiased approach to therapeutic development. Thus, rather than using rational design to select a target and develop a targeting moiety, cycles of <u>S</u>ystematic <u>E</u>volution of <u>L</u>igands by <u>Ex</u>ponential Enrichment (SELEX) are employed in cell culture or *in vivo* to identify aptamers against unknown targets. Antibody drug conjugates (ADCs) have shown some efficacy for GBM but are limited by their large size and thus depend on leakiness of the BBB. We have recently applied *in vivo* SELEX to develop anti-GBM aptamers (six-fold smaller in mass than IgG antibodies) and to select aptamer-drug conjugates. Here we report attempts to focus aptamer selection toward internalizing drug-delivery targets and resulting challenges involving loss of tumor specificity *in vivo*.

**GRAPHICAL ABSTRACT:** 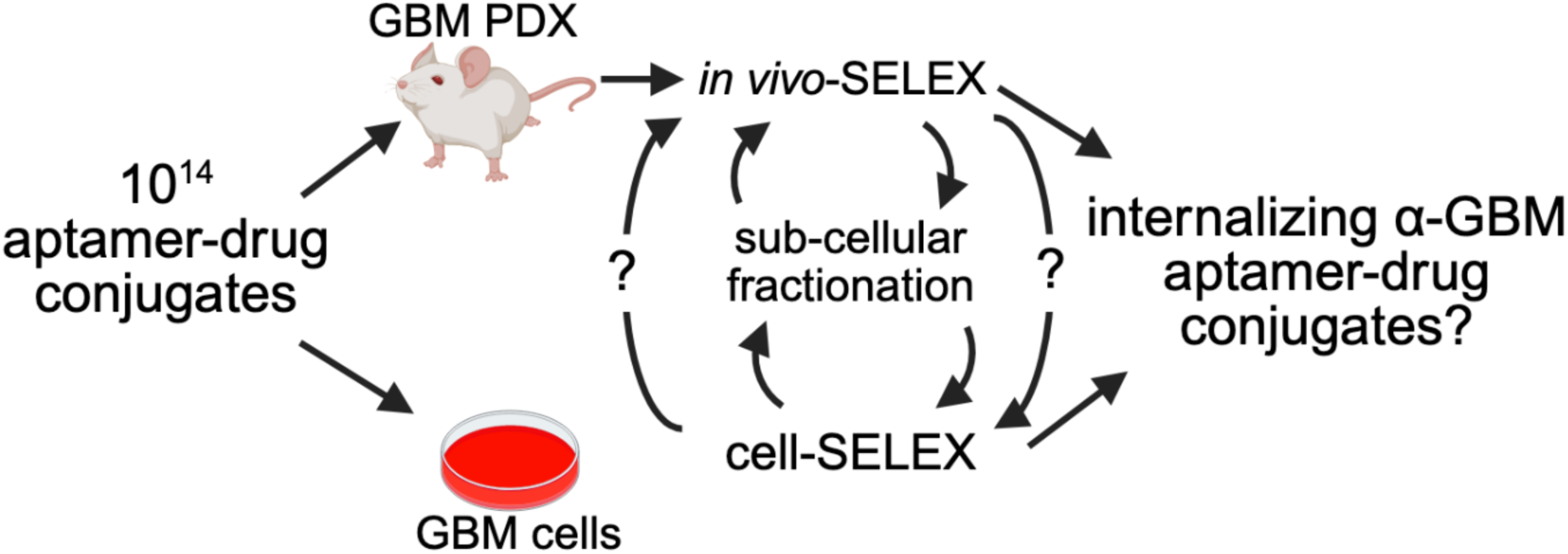

## INTRODUCTION

Glioblastoma (GBM) is the most common primary malignant brain tumor. Unfortunately, GBM has an exceptionally poor prognosis, being nearly universally fatal (1). Despite the urgent need for therapeutics, GBM treatment has remained essentially unchanged for over two decades since temozolomide (TMZ) was approved by the United States Food and Drug Administration in 2005 (2, 3). Challenges to therapeutic development include the blood brain barrier (BBB), the diffuse and infiltrative growth pattern of the tumor, and marked intratumoral heterogeneity (4–6).

Antibody-drug conjugates (ADCs) have shown some promise against GBM in pre-clinical and early stage clinical trials, all have failed Phase 3 clinical trials as of current(7, 8). Though ADCs, such as Deputux-M, target antigens involved in receptor-mediated transcytosis (RMT) at the BBB, they were not optimized for RMT and rely on BBB leakiness(7). This is not a efficacious therapeutic strategy as BBB leakiness differs from patient-to-patient, by tumor location, and as a function of disease progression (7).

Aptamers are short synthetic, single strands of DNA or RNA or analogs that fold into three-dimensional shapes, analogous to antibodies, with potential for high target affinity and selectivity. As such, aptamer function arises from three-dimensional shape, not genetic coding in the nucleotide sequence. An 80-nucleotide (nt) aptamer has ∼six-fold lower mass than an IgG antibody, conferring increased potential to cross the BBB (9, 10). Previously, we demonstrated that *in vivo* SELEX utilizing an orthotopic patient derived xenograft mouse model could be used to successfully identified unconjugated and toxin-conjugated anti-GBM aptamers and aptamer-drug conjugates (ApDCs) (11, 12). The selected aptamers and ApDCs successfully home to orthotopic patient-derived GBM xenografts after i.p. injection in mice (11, 12). Beyond *in vivo* homing capability, these i*n vivo*-selected naked DNA aptamers show target cell binding *in vivo* and *in vitro*, tumor tissue staining, and pharmacokinetics measurable by quantitative PCR (qPCR) (11, 12).

However, this prior work also raised several key questions critical for refining *in vivo* SELEX methodologies to achieve clinical relevance. Challenges include i) how to effectively select tissue-homing, cell-specific, *internalizing* aptamers when toxin conjugates must reach intracellular destinations for activity; ii) whether toxin conjugation influences *in vivo* tissue homing, iii) whether tumor specificity can be achieved for ApDCs in a patient-derived xenograft (PDX) model whose BBB is more intact, and iv) whether *in vivo-*selected aptamers demonstrate enhanced *in vivo* homing relative to cell-selected aptamers that have never encountered a BBB before.

Here we address two main questions. We first directly evaluate whether aptamers trained by *in vivo*-SELEX demonstrate enhanced *in vivo* homing compared to aptamers trained by *in vitro* cell-SELEX. We then address whether focused *in vivo* selection demanding cell uptake into endocytic pathways preserves GBM homing and promotes toxicity of MMAE-conjugated DNA aptamers whose MMAE anti-microtubule toxin must be released by lysosomal cathepsin protease.

While, in principle, *in vivo-*SELEX should confer advantages, especially for an intracranial target, literature reports of successful *in vivo* performance of aptamers developed by cell-SELEX motivated this direct comparison in GBM orthotopic PDX mouse models (13, 14). We therefore undertook cell-SELEX experiments with independent DNA libraries trained with and without MMAE toxin conjugation. We demonstrate that GBM cell binding of the top three resulting aptamer candidates from cell-SELEX libraries with or without MMAE bound target cells *in vitro* significantly more than a negative control, independent of their MMAE conjugation status, implying tolerance to MMAE conjugation. This result differs from what we had previously observed for *in vivo*-selected DNA aptamers (12). Importantly, we show that, unlike *in vivo*-selected DNA aptamers, these cell-selected aptamers do not home to target GBM PDX cells *in vivo*. Rather, we detect significant accumulation of these aptamer-MMAE conjugates and unconjugated aptamers in the lung, regardless of MMAE conjugation. Further, for ApDCs capable of GBM cell binding *in vitro*, no reproducible toxicity was detected *in vitro*. As in our previous *in vivo* selections, we hypothesized that this lack of ApDC toxicity was due to failure of ApDCs to bind internalizing GBM cell surface receptors, so necessary delivery of conjugates to activating lysosomal proteases fails.

We therefore describe SELEX attempts involving *in vivo* SELEX or hybrid (combined cell-SELEX and *in vivo*-SELEX) in the same orthotopic PDX model, but rewarding aptamers not only for GBM homing but also for subcellular localization in the endolysosomal pathway. Importantly, evaluation of the biodistribution of the resulting libraries and their leading candidates *in vivo* shows loss of GBM homing. Thus, future *in vivo* SELEX strategies will be needed to simultaneously optimize ApDC target cell binding, cell internalization, and toxin release.

## MATERIAL AND METHODS

### PDX models

Short-term PDX explants were obtained from the Mayo Clinic Brain Tumor Patient-Derived Xenograft National Resource. PDX models were established directly from patient biopsy or surgical samples as flank tumors in athymic nude mice and maintained as previously described (19). These tumors were then used to establish short-term explant cultures, intracranial (orthotopic) xenografts, or subcutaneous xenografts (19). Each PDX model is highly annotated with multi-omic characterization and relevant clinical data as previously described (20). G43 was selected as a highly aggressive PDX originating in a 69-year-old male patient with grade IV IDH-wild type GBM in the right temporal lobe.

### Cell culture

GBM PDX lines were maintained as previously described (15). Short-term explant cultures of G43 and G43-eGFP were grown in DMEM (Fisher Scientific #MT10013CV), supplemented with 10% fetal bovine serum (Gibco #220346) and 1% penicillin/streptomycin (Life Technologies #221674) at 37°C, in room air supplemented to 5% CO_2_. Identities of G43 and G43-eGFP were verified by short tandem repeat analysis. PDX lines were tested every 6 months for mycoplasma using a MycoAlert mycoplasma detection kit (Lonza #LT07-418). SVG-A were grown in DMEM (Fisher Scientific #MT10013CV), supplemented with 10% fetal bovine serum (Gibco #220346) and 1% penicillin/streptomycin (Life Technologies #221674) at 37°C, 5% CO_2_.

### Random DNA library synthesis for cell SELEX

Details of DNA library synthesis methods are provided in Supplementary Data available at NAR online.

### Orthogonal Cell SELEX

Details of SELEX methods are provided in Supplementary Data available at NAR online.

### *In vivo* SELEX in orthotopic PDX mice

Details of SELEX methods are provided in Supplementary Data available at NAR online.

### Synthesis of individual candidate ApDCs

Individual candidate aptamers were synthesized by IDT (Coralville, IA). ApDCs were then created in-house via PCR using an MMAE-conjugated upper primer, scaling up the conventional 100 µL reaction (10 μL 10× *Taq* DNA polymerase buffer, 10 μL 10× 1 mg/mL BSA, 8 μL 50 mm MgCl_2_, 8 μL 2.5 mm dNTPs, 10 μL library, 10 μL 5 μM 5’MMAE forward primer, 10 μL 5 μM 5’phosphorylated reverse primer, 32.6 μL water, 1 µL of 100 µM template and 1.4 μL *Taq* DNA polymerase) as needed to 1-4 mL. Following thermal cycling, PCR reactions were pooled, DNA precipitated from ethanol and resuspended in 88 µL water supplemented with 10 µL 10× phage lambda exonuclease buffer and 2 µL of phage lambda exonuclease (New England Biolabs). The solution was incubated at 37°C for 30 min to hydrolyze the 5’-phosphorylated reverse strand. An equal volume of formamide was added to the reaction followed by heating at 90°C for 5 min. ApDCs were purified by electrophoresis through a 7.5% denaturing polyacrylamide gel at 600 V/23 cm for 2.5 h, visualized by UV shadowing, and a gel fragment excised with a razor blade. The gel fragment was minced and ApDCs isolated by incubation in 0.3M NaOAc as described in Supplementary Methods.

### Cell association assays

For cell association experiments, explants of the parental G43 or G43-eGFP-fLUC2 modified cells were used. Cells were plated overnight in 24-well plates (Corning #3512) such that cultures were ∼80% confluent on the day of the experiment. Cells were washed with wash buffer (cell-selections: PBS containing 5 mM MgCl_2_ and 4.5 g/L glucose; *in vivo-*selections: PBS containing 1 mM MgCl_2_ and 4.5 g/L glucose) prior to adding heated and snap-cooled aptamer or ApDC to 50 nM final concentration in selection buffer (cell-selections: 5 mM MgCl_2_; *in vivo*-selections: 1 mM MgCl_2_ in PBS) with 4.5 g/L glucose, 0.1 mg/mL competitor nucleic acid [cell-selections: tRNA (Sigma Aldrich, #R8759-2KU); *in vivo*-selections: sheared salmon sperm DNA (Invitrogen #15632011V)] and 0.1 mg/mL BSA (New England BioLabs #B9200S). Cells were incubated with 50 nM aptamers or ApDCs for 30 min at 37°C. Cells were then washed three times with the corresponding wash buffer.

For qPCR, cells were recovered by scraping on ice in 100 µL wash buffer. Wells were washed once with a second portion of 100 µL wash buffer to collect any remaining cells. Collected cells were subjected to centrifugation at 500 × g for 5 min and the supernatant aspirated. Equal volumes of lysis buffer (50 mM Tris HCl, pH 7.4, 150 mM NaCl, 1% NP-40, 0.5% Na Deoxycholate, 1 mM EGTA, and 1 mM NaF) and Qiagen Plasmid Extraction Kit Buffer P1 were added to the cells, with vortex mixing to resuspend. The lysate was incubated on ice for 30 min prior to sonicating each sample for 10 s at power 8 (Sonic Dismembrator 60, Fisher Scientific). Lysates were heated at 95°C for 10 min and then subjected to centrifugation for 5 min at 13,100 × g in a microcentrifuge. Samples of the resulting supernatant were used for qPCR (QuantaBio PerfeCTa #95071-012). Aptamers were either quantified by their specific standard curve or it was validated that they had similar standard curves and one aptamer could be used as a representative curve for all. Protein was quantified by Qubit Protein Assay Kit (Invitrogen #Q33211).

### Western Blot Analysis

The following primary antibodies were used: anti-GAPDH (Abcam, #ab8245), anti-β-Tubulin (Invitrogen, #41-4510-80), anti-cathepsin B (Proteintech, 12216-1-AP), anti-LRP6 (C5C7) Cell Signaling, #2560), anti-GM130 (D6B1) (Cell Signaling, #12480), anti-Rab5 (C8B1) (Cell Signaling, #3547), anti-LAMP1 (Abcam, #ab24170), anti-SDHB (Abcam, #ab14714), anti-Rab7 (Abcam, #ab50533), anti-Rab9 (Abcam, #ab179815), anti-Lamin A/C (Santa Cruz, #sc-7292), and anti-Na^+^/K^+^ ATPase (Sigma-Aldrich, #05-369-25UG). Primary antibody binding was detected using a goat anti-rabbit conjugated IRDye 680RD (Li-Cor, #926-68071) or a goat anti-mouse conjugated IRDye 800CW (Li-Cor, #926-32210).

### Analysis of individual aptamer biodistributions

Mice were stereotactically injected with 10^5^ parental G43 cells without GFP expression (details in Supplementary Methods). Mice were closely monitored for weight and behavioral changes as signs of tumor progression. Animals approaching moribund status were enrolled in the study. Aptamer cocktail solutions (400 pmol/400 μL; 50 pmol of each of 8 aptamers or ApDCs) were created in PBS containing 5 mM MgCl_2_, heated for 5 min at 90°C and snap-cooled on ice for 10 min. Control vehicle solutions lacking aptamers were created at the same time. Mice were injected i.p. with 400 μL aptamer cocktail (400 pmol aptamer total) or vehicle. 30 min post-injection, mice were euthanized, subjected to scrupulous transcardial perfusion (see Supplementary Methods) and the brain, heart, lungs, spleen, liver, kidneys, bone marrow (extracted from both femurs by centrifugation at ≥10,000 × g), and sciatic nerve were collected in tared tubes and immediately frozen on dry ice and stored at -80° C. Toluidine blue staining facilitated GBM tumor resection.

Subsequent brain dissection involved warming the brain to -20° C and placing it in a dissection block maintained at -20° C. Fresh razor blades were placed as indicated in **Supplementary Figure S2F**. Two 1-mm brain sections were obtained and stained with toluidine blue (0.33 mg/mL) for 30 s. Excess toluidine blue was removed from the tissue with a pipette and the brain was dissected away from the tumor using differential toluidine blue staining of tumor tissue as a guide (**Supplementary Figure S2**). After sectioning, the remaining posterior tumor was dissected from the right hemisphere. The cerebellum was isolated.

For biodistribution experiments, an optimized aptamer extraction method was used as previously described (11, 12). The lysate was analyzed by qPCR (QuantaBio PerfeCTa #95071-012) using sequence-specific primers (**Supplementary Table S1**).

### Toxicity analysis

A CellTiter-Glo assay (Promega #G7571) was used to determine aptamer toxicity on cultured GBM39 target cells. Aptamers were mixed with complete media and added to target cells. Six days post-treatment, a CellTiter-Glo assay was performed on triplicate samples according to the manufacturer’s protocol.

## RESULTS

### ApDCs selected by cell-SELEX using independent DNA libraries +/-MMAE conjugation

We sought to evaluate whether aptamer selection with or without MMAE toxin conjugation had any impact on post-SELEX modifications. In addition to this question was the desire, to better understand the benefits and limitations of *in vivo-*SELEX vs. cell-SELEX for anti-GBM aptamer, which require BBB penetration. Thus, we performed two parallel *in vitro* cell selections with independent DNA libraries. We refer to independent libraries as “orthogonal” if they require different reverse primers for PCR amplification (our standard procedure) to limit cross-contamination during processing. For cell-SELEX, aptamer or ApDC libraries were incubated with normal human astrocytes (SVG-A cells: negative selection) for 30 min. Unbound sequences were then transferred to cultured explants of GBM PDX (G43 cells: positive selection) for 30 min prior to washing and recovery of aptamer or ApDCs bound to the target cells (**Figure 1A**). The -MMAE (**Figure 1B**) and +MMAE (**Figure 1C**) libraries completed ten SELEX rounds to determine how selection with or without toxin affected tolerance to toxin conjugation post-selection. After 10 rounds of cell-SELEX, libraries underwent deep sequencing and varying degrees of exponential enrichment were observed in both cases (**Figure 1D**). Interestingly, the top three +MMAE candidates, were about 4-fold more enriched than the top 3 -MMAE candidates. Neither library shared random regions of the top sequences. The top three candidates from both libraries were then individually synthesized and tested for target cell affinity *in vitro* with and without MMAE conjugation. Although different candidates varied in their abilities to tolerate toxin modification as evidenced by cell binding, all candidates demonstrated target cell specificity vs. negative control cells and negative control aptamer (7546 and 7547), regardless of MMAE conjugation (**Figure 1E**).

**Figure 1.**
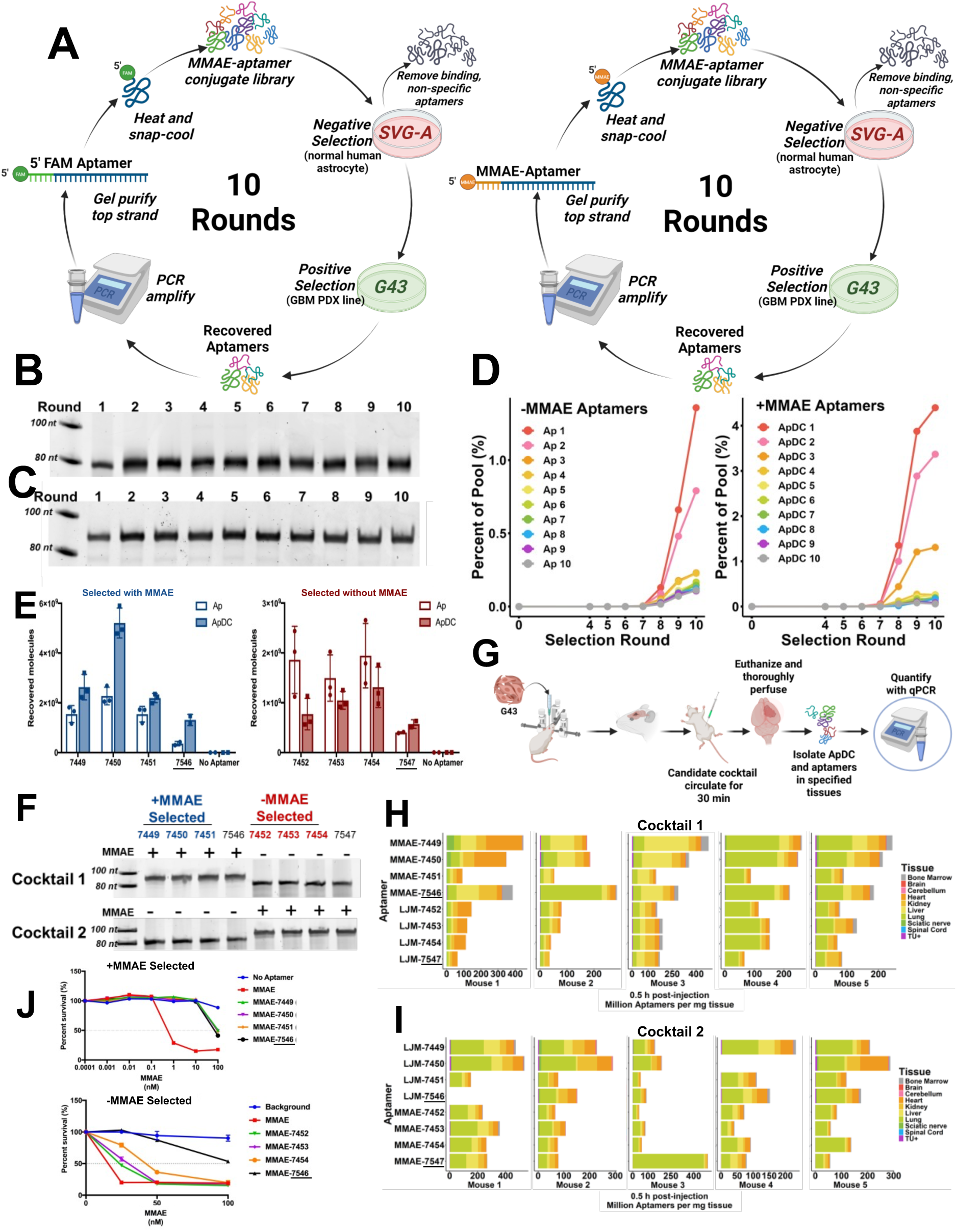
Orthogonal cell SELEX experiments utilizing libraries +/-MMAE conjugation. A. Schematic diagram of parallel selections. B-C. Confirmation of library length for either unconjugated (B) or conjugated (C) libraries over 10 rounds of cell SELEX. D. Deep sequencing results for the top 10 aptamer or ApDC candidates from the +/-MMAE libraries after 10 rounds of cell SELEX. **Supplementary Table S1** translates the generic aptamer names to the numerical serial code. E. Cell association of the top three candidates from each of the +/-MMAE libraries relative to respective negative controls (7546 and 7547). F. Schematic diagram of the experiment evaluating the biodistribution of cell-selected candidates in mice with orthotopic PDX brain tumors (F -I). Cell-SELEX candidate aptamers were evaluated for their respective biodistributions +/-MMAE. Two cocktails were created: Cocktail 1 contained selected versions of the candidates in the conjugation state used during selection and Cocktail 2 contained the opposite conjugation state of the same candidates (G). Mice received an i.p. injection of 400 µL (50 pmol/candidate) of either cocktail. After 30 min mice were euthanized and thoroughly perfused. Individual aptamers were quantified in the indicated tissues by qPCR with aptamer-specific primers (H-I).

In light of data from prior *in vivo* selections in our lab (12), this finding is striking as it indicates that the inability to tolerate post-SELEX modifications may be especially problematic for aptamers selected *in vivo* vs. *in vitro*.

### Biodistribution of cell-selected aptamers +/-MMAE conjugation

We used qPCR to determine the *in vivo* biodistribution of cell-selected aptamers that bound target cells *in vitro* +/-MMAE conjugation. We sought to answer two questions: (1) is *in vivo* SELEX important to identify ApDC candidates with target specificity *in vivo*, and (2) does conjugation to MMAE change the biodistribution of ApDCs relative to unconjugated aptamers. Two cocktails of the top three candidates and negative control from both +/-MMAE libraries were created: Aptamers in Cocktail 1 were injected in the conjugation state originally selected while aptamers in Cocktail 2 were synthesized in the opposite conjugation state (**Figure 1F**). As illustrated in **Figure 1G**, to determine the biodistribution of the cell-selected aptamers and ApDCs, cocktails were injected i.p. into mice bearing an orthotopic G43 tumor. 30 min later mice were euthanized and scrupulously perfused. Organs were collected and aptamers were extracted and quantified by qPCR with aptamer-specific primers. Interestingly, at 30 min the biodistribution appeared to reflect the degree of physiological tissue vascularity more than target homing. There was no difference between MMAE-conjugated and -unconjugated molecules (**Figure 1H-I**). Strikingly, homing to the orthotopic brain tumor previously observed for unconjugated in vivo-selected aptamers was lost (11). A very small fraction of only a few aptamers was detectable in the PDX GBM. This result appeared to be mouse-dependent rather than sequence-dependent. It is possible that the leakiness of the BBB in individual tumors may influence GBM uptake. Furthermore, ApDC conjugates from the +MMAE library (7449 and 7450) displayed slightly more tumor homing than any sequence from the -MMAE library, regardless of original conjugation state. Notably, the scrambled negative control sequence for the +MMAE library, 7546, associated with the GBM as well as the two selected anti-GBM aptamer candidates in every mouse, indicating that the scant tumor uptake is nonspecific. The large variability between mice in this experiment prevented meaningful statistical deduction. We were left with the finding that there was no difference in organ accumulation between +/-MMAE versions of aptamers selected against GBM cells *in vitro* (**Supplementary Figure S3**). Thus, 30 min post-injection, no evidence was found that MMAE conjugation caused differential accumulation in highly-vascularized organs such as lung. Further work would be required to determine results at later timepoints.

Although our previous *in vivo-*selected candidates (11, 12) had been selected and tested in a different GBM PDX model (G39) we compared the accumulation of these *in vivo*-selected aptamers in brain, lungs, and respective tumor at 30 min vs. the current cell-selected anti-G43 candidates at the same timepoint (**Supplementary Figure S4**). It is important to note, this is not a perfect comparison as the *in vivo-*selected candidates were intravenously (i.v.) injected and the cell-selected were intraperitoneal (i.p.) injected, but we have previously shown that tumor accumulation for these *in vivo*-selected candidates (ApDCs 2 and 5) is the same regardless of injection route and off-target tissue accumulation was slightly more with i.v. injection. Interestingly, accumulation of cell- and *in vivo*-selected aptamers in off target tissue such as the lung was similar at 30 min post-injection. However, aptamer accumulation in the GBM tumor was strikingly different. Aptamer accumulation in lung for *in vivo-*selected candidates was not different from cell-selected candidates (regardless of MMAE conjugation state). More importantly, accumulation in the PDX GBM tumor by the *in vivo-*selected candidates was ∼15-fold higher relative to cell-selected candidates. Furthermore, if we take into account the difference in the leakiness of the BBB of the two different models and select a later timepoint, such as 2h, where the negative control scrApDC from the *in vivo* selection cohort is comparable to the cell-SELEX negative controls 7546, and 7547, the *in vivo* selected candidates are still ∼5-fold higher in the tumor relative to the cell-selected candidates. Additionally, when evaluating the cell-SELEX candidates in isolation, there was no difference in GBM delivery between the cell-selected candidates and any of the three negative controls. *Taken together, these results strongly suggest that in vivo SELEX is necessary to identify aptamers with strong GBM specificity in vivo*. Our results also suggest that lung accumulation may reflect tissue vascularity and/or the challenge of incomplete tissue perfusion.

Given that successful *in vivo* homing of cell-selected aptamers has been reported in the literature for some tissues other than GBM, we sought to determine if the lack of homing of our cell-selected aptamers to the orthotopic GBM might be due to the BBB. We repeated a biodistribution experiment, but this time used mice bearing G43 flank tumors where a BBB is not present (**Supplementary Figure S5A**). We hypothesized that eliminating the need to cross the BBB would facilitate accumulation of aptamers/ApDCs in the flank tumors. In fact, only a slight increase of flank tumor uptake was observed for all sequences in this location (**Supplementary Figure S5B**). No differences in biodistribution pattern were observed between aptamers +/-MMAE. We conclude that the failure of cell-selected aptamers to home to the G43 tumors (orthotopic or flank) is independent of the BBB. The lack of significant improvement in GBM homing for flank tumors could be due to a number of factors including the inability of the cell-selected aptamers to extravasate from blood vessels into the target tissue and/or the target ligands on cultured GBM cells differing from those *in vivo*.

### Lack of toxicity of cell-selected ApDCs

We tested the *in vitro* cell culture toxicity of cell-selected MMAE-conjugated candidates from both +/-MMAE libraries (**Figure 1J**). While the cell-selected aptamers were trained in a simpler context (cultured cells) and bind cultured GBM cells more than a negative control aptamer, reproducible toxicity in a six-day assay was not observed for any of the MMAE conjugates, relative to a negative control MMAE conjugate. This result suggested to us that dominant cell-selected aptamers bind non-internalizing targets on GBM cells. This, in turn, suggested that the SELEX strategy should be modified to reward uptake into the sub-cellular lysosomal pathway where protease (e.g. cathepsin) release of the MMAE toxin is believed to occur.

In summary, studies with cell-selected aptamers confirmed that i) *in vivo* selection is essential to select aptamers with strong GBM-homing properties *in vivo*; ii) top sequences from a cell-SELEX library do not demonstrate GBM cell toxicity in culture; and iii) some cell-selected aptamers tolerate post-SELEX toxin conjugation without biodistribution changes, though *in vivo* target specificity for these aptamers is unimpressive. Effects at later timepoints must be examined in future studies.

### A SELEX strategy to reward ApDCs that enter an intracellular lysosomal pathway

With the goal of developing functional, anti-GBM homing ApDCs, we returned to *in vivo-*SELEX in the same G43 model used for cell selections, but introduced a protocol intended to reward toxin delivery to the endolysosomal pathway. We maintained the conventional MMAE toxin and valine-citrulline (VC) p-aminobenzylcarbamate (PAB) linker, in which toxin release requires cleavage of ApDCs by the lysosomal enzyme, Cathepsin B. In pilot studies designed to isolate molecules that were internalized and cleaved by Cathepsin B, we tested but ruled out two potential physical selection strategies. The first would exploit EDC-NHS coupling chemistry to capture 5’carboxyl-containing aptamers (the products of MMAE cleavage) (**Supplementary Figures S6-7**). The second would exploit the electrophoretic gel shift to higher mobility for ApDCs from which MMAE had been cleaved (**Supplementary Figure S8**). These approaches proved challenging because i) EDC-NHS coupling gives poor yields for trace concentrations of aptamers and ii) gel mobility shift-based aptamer purification was too inefficient for *in vivo* SELEX.

We then devised a cell fractionation protocol to enrich intact intracellular vesicles in the endolysomal pathway (**Supplementary Figures S9**). This approach relies on an endolysosome enrichment kit to isolate and amplify only aptamers more likely to deliver the MMAE toxin intracellularly. The goal was to overcome the demonstrated lack of toxicity of ApDCs relative to negative controls, which we had failed to achieve with our previous *in vivo* selections (**Supplementary Figures S10**) (11, 12). A slight improvement in *in vitro* toxicity was observed for candidates with the largest fold change between the input and the library recovered from one round of endolysosomal enrichment from cell culture (**Supplementary Figure S9**). This provided encouragement that the enrichment protocol might reward ApDCs that successfully deliver toxins intracellularly.

### *In vivo*-SELEX using an orthotopic PDX GBM model to select ApDCs that home to GBM and enter the endolysosomal pathway

The goal of the resulting *in vivo-*SELEX was to reward not only GBM homing *in vivo*, but also endolysosomal pathway localization such that the valine-citrulline peptide toxin linker would be more likely to be cleaved, releasing the toxic MMAE microtubule poison. Several parameters were changed in this *in vivo* SELEX attempt (**Supplementary Figure S11A**). To facilitate comparison with our prior cell-SELEX aptamer/ApDC experiments, we employed the same PDX line (G43) as was used for cell-SELEX, a line believed to give rise to orthotopic GBM tumors characterized by a more intact BBB than the previous model used for *in vivo*-SELEX (G39; **Supplementary Figure S11B**). We shortened the incubation time from 4 h to 30 or 90 min with the goal capturing aptamers in the process of internalization into the endolysosomal pathway, but prior to destruction in the lysosome. The time was selected based on cell culture estimates of fluorescent transferrin accumulation in G43 cells by endocytosis (**Supplementary Figure S11C**). In our previous selections, we rewarded any molecule that accumulated in the tumor after stringent perfusion 4 h post-injection. In contrast, we now applied sub-cellular fractionation to select aptamers found in the enriched endolysosomal extract of explanted GBM tumors after stringent perfusion at 30 or 90 min after i.p. injection.

**Figure 2.**
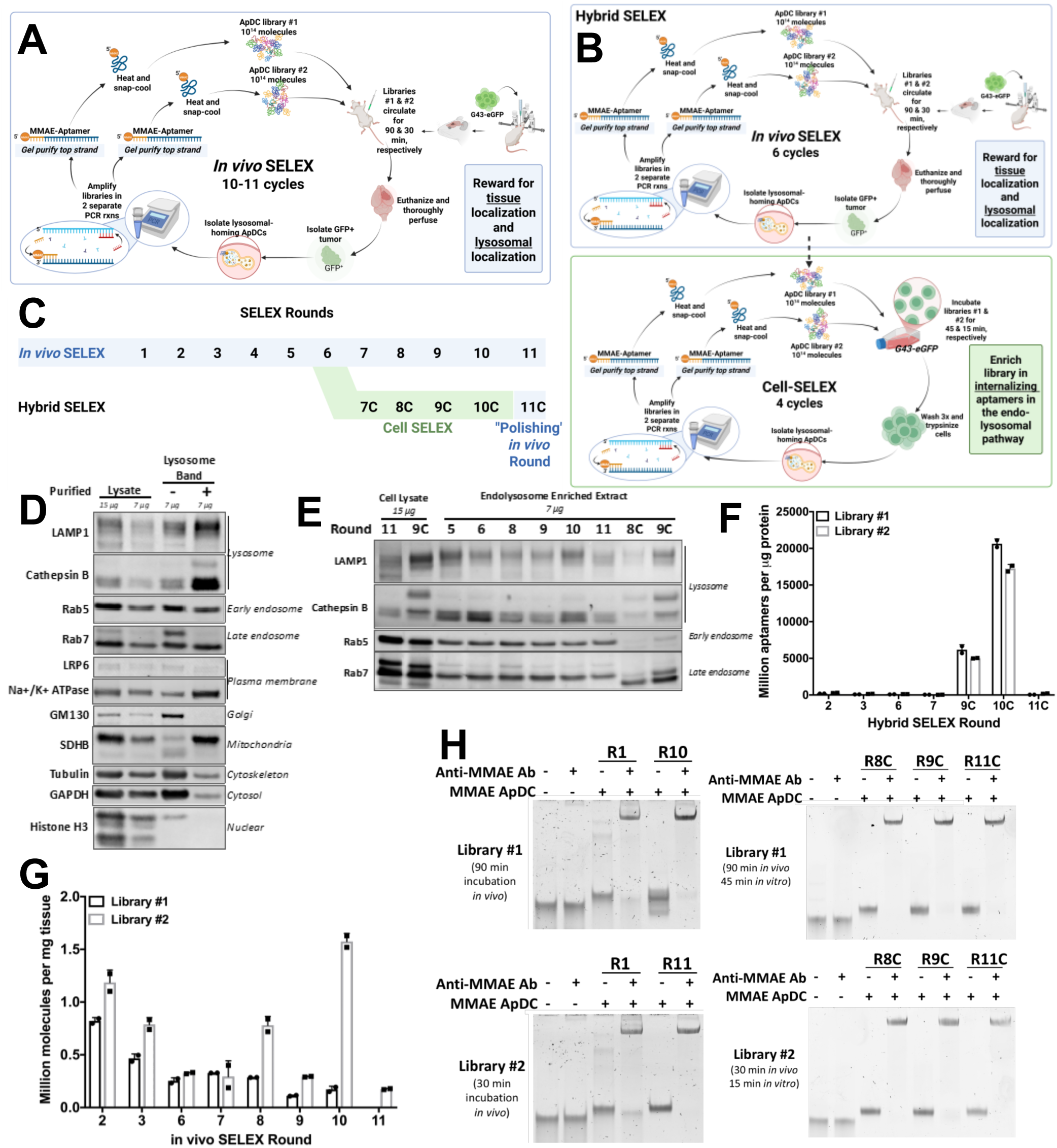
Multiplexed *in vivo* and hybrid SELEX experiments performed with orthogonal libraries and an orthotopic PDX model. (A-B). Schematic diagrams of the *in vivo* SELEX (A) and the Hybrid SELEX (B). C. Diagram outlining selection strategies. D. Western blot of example endolysosomal fraction from *in vivo* SELEX. E. Western blot of recovered endolysosomal samples from explanted GBM tumors at the indicated rounds of *in vivo* SELEX. (F-G). Aptamers recovered from the endolysosomal fractions as determined by qPCR. H. Confirmation of MMAE conjugation to the regenerated libraries was achieved with a gel mobility shift assay utilizing an anti-MMAE antibody and detection in a native polyacrylamide gel stained with SYBR gold.

A consequence of this change in reward strategy is that we sought to amplify only aptamers that quickly enter the degradation pathway, rather than molecules that escape degradation and accumulate in the tumor over longer times.

Not knowing ideal timing for this demanding goal, we employed two orthogonal libraries with different exposure times: a long-exposure library (90 min; Library #1) and a short-exposure library (30 min; Library #2). This was achieved by multiplexing each mouse at each SELEX cycle using sequential timed library i.p. injections (**Figure 2A**). Furthermore, we wished to evaluate whether the results of this selection would be improved using a hybrid SELEX strategy where the library was first trained by this method *in vivo* (6 rounds), then trained in cell culture (4 rounds), and then subjected to one final selection round *in vivo* (**Figure 2B-C**). For every round, after library incubation in the mouse, tissue was immediately collected and the crude endolysosomal fraction kit-purified for selective aptamer extraction. We confirmed that the resulting rapid sub-cellular extract fraction was enriched in early and late endosome and lysosomal markers, including Cathepsin B (**Figure 2D-E, Supplementary Figure S12**). Some contaminating mitochondria, and cytoskeletal, cytosolic, and membrane proteins were detectable in this crude fraction, while nuclear proteins were excluded. For Round 1, the recovered sample was added to a PCR reaction containing both primer sets for several rounds of amplification to create multiple copies of every sequence. For all subsequent rounds, the recovered sample was divided between the two PCR reactions and amplified to the optimal number of cycles. Both PCR reactions utilized a 5’MMAE-conjugated forward primer, which we had previously shown to be PCR-compatible (12), thus restoring the conjugation of the library to the toxin each round. After 10-11 rounds of *in vivo*-SELEX using this method, aptamers recovered from each round were quantified by qPCR (**Figure 2F-G**). Interestingly, only the cell-SELEX rounds from the hybrid selection demonstrated convincing exponential enrichment, while the protocol involving only *in vivo* SELEX lacked convincing exponential enrichment, unusual for *in vivo* selections in our experience. Care was taken to confirm that both libraries in the hybrid and *in vivo-*only selection protocols maintained toxin conjugation throughout the selection process (**Figure 2H, Supplementary Figure S12D**).

### Aptamer biodistribution determined by deep sequencing after selection for GBM endolysosomal delivery

After the final round of selection, all organs were collected from the mouse and processed for deep sequencing to determine the biodistribution of MMAE-conjugated aptamer candidates. The GBM brain tumor for this round was divided in half: one half was processed immediately for endolysosomal fractionation and the other half was flash frozen and processed as a bulk tissue with other organs to quantitate aptamer delivery per mg tissue as in our previous work (**Figure 3A**).

Final libraries were initially deep sequenced by Azenta’s AmpliconEZ Seq approach to confirm enrichment. Strikingly, there was a difference in nucleotide distribution between the two libraries (**Supplementary Figure S13A-B**). Library #1 (long-exposure) gave rise to guanine-rich candidates suggestive of potential guanine quartet structures. This library was difficult to amplify by PCR. Selected aptamers from Library #2 (short-exposure) were thymine-rich, as had been observed in our previous *in vivo* selections (12).

We assumed that sequences enriched in the bulk tumor preparation would be comparable to the endolysosomal preparation, so we first analyzed the top sequences found in the bulk tumor at the final SELEX round (**Figure 3**). The top 10 such sequences were evaluated for enrichment in the libraries over the prior rounds of selection. However, with the exception of one case, exponential enrichment was not observed in either library (**Figure 3B-C**). The top 10 sequences in both libraries were then evaluated for biodistribution in the final mouse, for both the *in vivo-*only selection (**Figure 3D-E**) and the hybrid selection (**Figure 3F-G**). Strikingly, the impressive GBM tumor homing observed in our previous *in vivo* selection with unconjugated and unconjugated aptamers selected only for GBM retention (11, 12) was completely lost. Indeed, when the recovered libraries were evaluated for fraction retained in the GBM tumor relative to the sum of other organs at the time of harvest, and compared to the cell-selected candidates, no difference was observed. Certain sequences could be found with higher abundance in the GBM, but these sequences represented only a small fraction of the pool and were not detected in other organs (**Supplementary Figure S14**). We conclude that the endolysosomal selection strategy rewards an uptake process that is not GBM-specific.

**Figure 3.**
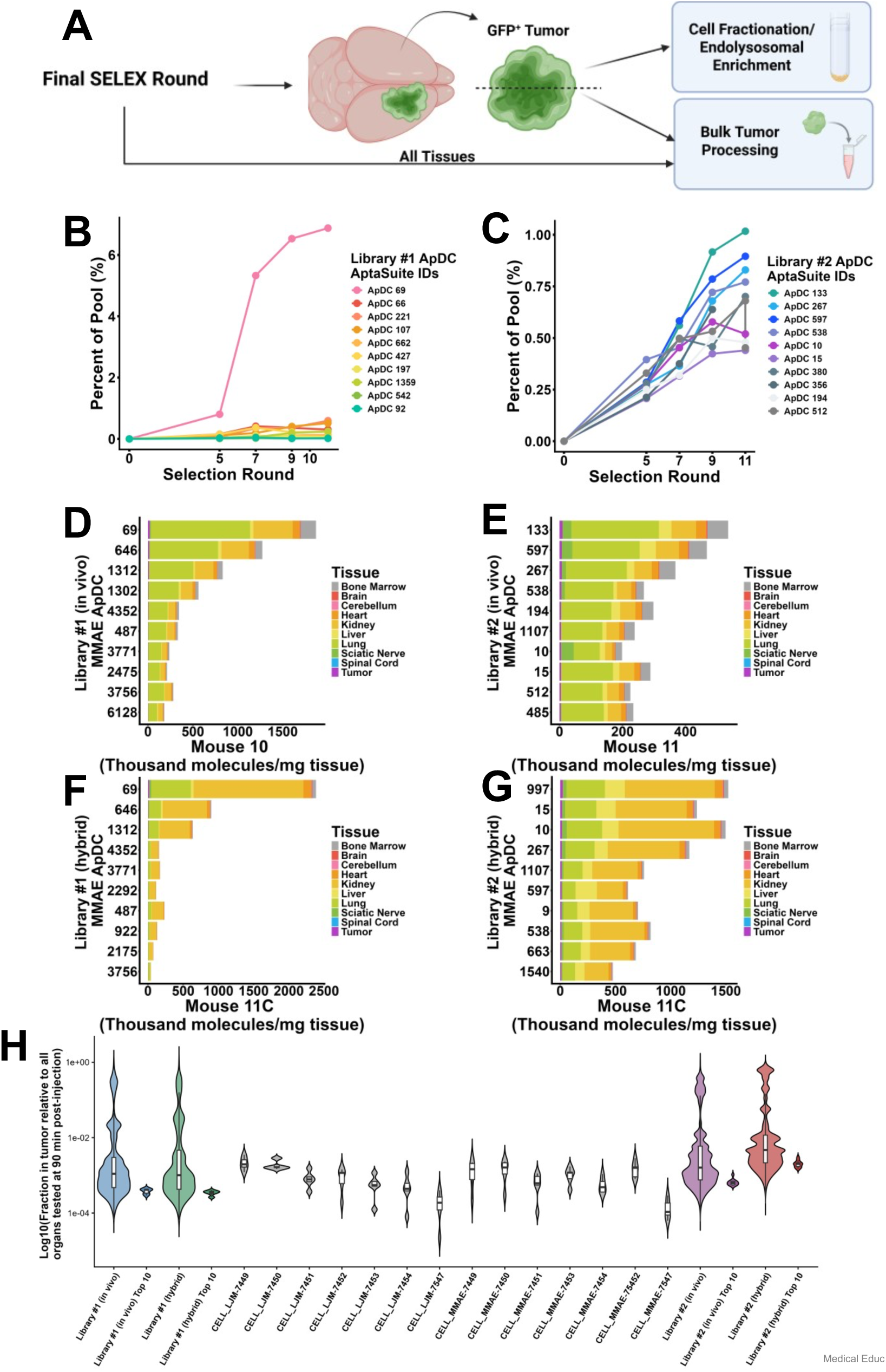
Biodistribution of top candidates from bulk tumor as determined by deep sequencing after the final round of *in vivo* SELEX. A. Schematic diagram of tumor division after the final round of SELEX. The sample that underwent cell fractionation was used to restore the library after the final round. (B-C) Enrichment of top 10 sequences in the tumor from *in vivo-*only SELEX in the libraries over the course of the selection. (D-G) Biodistribution of the top 10 sequences found in the bulk tumor in the *in vivo-*SELEX only library mouse (D-E) and the Hybrid SELEX library mouse (F-G). H. Comparison of the four libraries that resulted from *in vivo* and hybrid SELEX by evaluating Log_10_ of the fraction of an aptamer in the tumor relative to all organs tested at the time of harvest, and comparing them to the values achieved by the +/-MMAE cell-selected aptamers.

We evaluated results for top individual aptamer sequences in the library from the endolysosomal fraction (**Figure 4**). Strikingly, the top 10 sequences in long-exposure Library #1 demonstrated exponential enrichment, suggesting that the selection was successful. The top sequences in short-exposure Library #2 increased in a more linear manner (**Figure 4A-B**). Seeking to better understand this difference, we evaluated the correlation between the sequences in the bulk tumor and the sequences in the endolysosome fraction (**Figure 4C-D**). Selected aptamers from long-exposure Library #1 showed a poor correlation between representation in the bulk tumor and in the endolysosomal fraction, whereas selected candidates from short-exposure Library #2 demonstrated a high correlation. We evaluated bulk tumor processing to determine if the process depleted the library of the aptamers enriched in the endolysosomal fraction, but did not detect a significant loss of these markers (**Supplementary Figure S15**).

**Figure 4.**
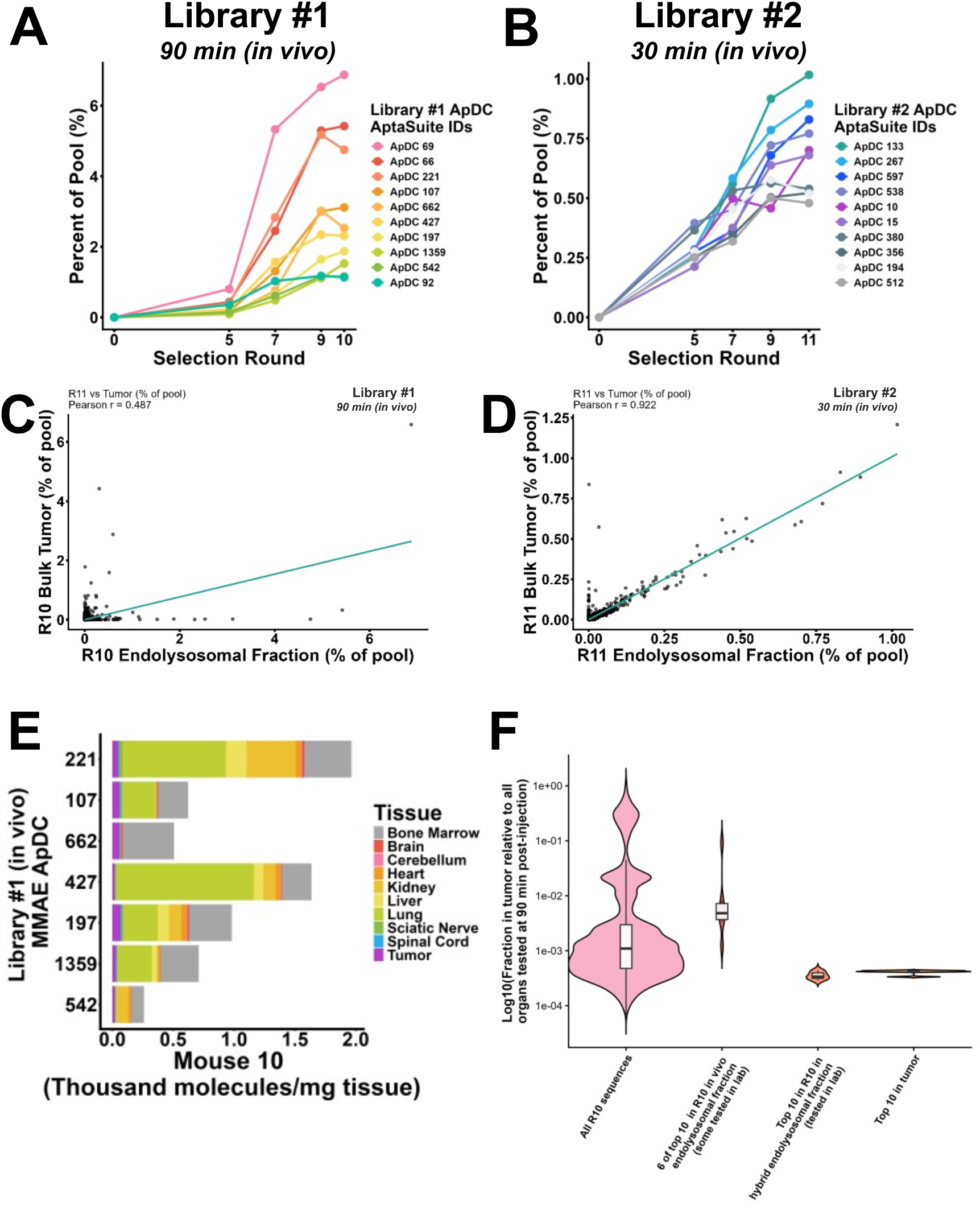
Enrichment and biodistribution of top candidates from the endolysosomal fraction as determined by deep sequencing after the final round of SELEX. (A-B) Enrichment of top 10 sequences in the endolysosomal fraction from *in vivo-*only SELEX in the libraries over the selection. (C-D) Correlation plots for the sequences enriched in the bulk tumor vs. the endolysosomal fraction in Libraries #1 (C) and #2 (D). E. Biodistribution of seven of the top sequences enriched in the endolysosomal fraction. F. Comparison of Library #1, the top endolysosomal fraction sequences from the *in vivo-*only SELEX, the top endolysosomal fraction sequences from the hybrid SELEX, and the top 10 in the tumor from the *in vivo-*only SELEX evaluating Log_10_ of the fraction of an aptamer in the tumor relative to all organs tested at the time of harvest.

We noted that the biodistribution of the top sequences in the endolysosomal fraction at the final round for long-exposure Library #1 had significantly improved tumor homing relative to all other sequences. This was only observed for the ten most enriched aptamers from the *in vivo-*only SELEX, not from the hybrid SELEX (**Figure 4E-F**, **Supplementary Figure S17A**). We hypothesize that the later timepoint reduced nonspecific aptamer signal sufficiently for the desired selection to occur. The weakness of this hypothesis is that when all four libraries were compared, aptamers in short-exposure Library #2 from the hybrid SELEX protocol demonstrate the highest fraction in the GBM relative to all other organs at the time of harvest (**Supplementary Figure S17B**). As noted, short-exposure Library #2 demonstrated a good correlation between the aptamers in the bulk tumor and in the endolysosomal fraction. We conclude that the endolysosomal fraction is contaminated by some cell surface-bound aptamers. It is thus likely that for short-exposure Library #2, endolysosomal enrichment was acting similarly to bulk tumor accumulation because contaminating aptamers had not yet been internalized. Enrichment may have simply been detecting all tumor-homing aptamers, including background in the imperfect endolysosomal fraction. At early times, such aptamers appear to be preferentially selected. At later times, more aptamer internalization has occurred, and the desired selection is possible. We therefore hypothesize that tumor homing might be improved at later times to further reduce background selection and further enhance tumor homing selection.

## DISCUSSION

Aptamers are developed through the principles of natural selection, permitting the use of an evolutionary strategy as opposed to a rational design strategy for therapeutic development. Thus, the required mechanisms by which an aptamer enters the bloodstream from the peritoneal cavity, crosses the endothelial barrier, infiltrates a tissue, binds to a specific cell, internalizes into that cell, and homes to a specific subcellular organelle are deliberately not engineered prospectively. Rather, a selection strategy is designed to reward aptamers from vast random libraries for successfully homing to the target location after successful navigation through all of these barriers (and, presumably, many more). This evolutionary approach highlights aptamer selection as a promising strategy for addressing challenging diseases such as GBM, which urgently require fundamentally different approaches for effective therapeutic development.

However, the full potential of aptamer selection remains largely unrealized, as evidenced by the limited advancement of aptamers in clinical development and the challenges our experiments have revealed. Aptamers have typically been selected against proteins or cells and then assumed to be competent for *in vivo* homing. Prior to 2025, aptamers relevant to neuro-oncology were exclusively developed via protein- or cell-SELEX. It is important to note that many of these aptamer selection strategies were therefore based on rational target selection.

Our work is guided by the hypothesis that *in vivo* aptamer target tissue delivery can be achieved by the unbiased SELEX approach itself. A long-term goal is to discover novel delivery strategies for GBM by understanding the mechanisms of successful GBM-homing aptamers selected from hundreds of trillions of random candidates by *in vivo* SELEX. This contention remains to be proven, and systematic experiments are needed to optimize clinically-relevant aptamers. Although the first *in vivo* SELEX experiment was published in 2010, the past 16 years have seen only 10 reports of successful *in vivo* SELEX experiments, demonstrating that much remains to be optimized in this strategy.

We deliberately chose to use unmodified DNA aptamers in our work. This decision was practical. It is vastly more expensive and time-consuming to select and synthesize aptamer analogs as candidates. In preliminary efforts to reduce triggering of the cGAS-STING innate immunity pathway, we designed primers lacking CpG dinucleotides, but we do not know if this strategy is valuable, and no effort was made to avoid CpG dinucleotides in aptamer random regions. Nuclease-resistant analogs such as aptamers with 2’-fluoropyrimidine RNA backbones would undoubtedly increase aptamer lifetimes *in vivo*, but the present studies nonetheless serve as proof of concept. Additionally, we argue that faster degradation of unmodified DNA oligonucleotides may actually aid in decreasing selection background, which is critically important when selecting aptamers that home to the endolysosomal pathway in a brain tumor *in vivo*.

For this work we used the G43 GBM PDX model that grows well in culture and is typically characterized by a more intact BBB in orthotopic xenografts relative to the G39 GBM PDX model employed in our previous work (11, 12). This choice sought to illuminate if *in vivo* SELEX is effective for PDX tumors characterized by different degrees of BBB leakiness. Additionally, we chose to work with a GBM model that is currently un-targetable by ADC therapies because it does not overexpress EGFR (7).

In our *in vivo* SELEX design, we deliberately created a reward protocol employing GBM cells at early passage, providing the highest yield of material for orthotopic injection, and permitting efficient SELEX timing. The timing of tumor seeding by orthotopic injection was staggered to generate mice with tumors of appropriate size for *in vivo* SELEX every 3-4 days. A limitation of our reward strategy based on amplifying aptamers recovered from the endolysosomal sub-cellular fraction was that the rapid fractionation kit does not yield scrupulously pure material such that some aptamers not in the endolysosomal pathway were likely also amplified at each round. Mitochondria were the most obvious contaminant. The preparation kit removed significant cytosolic and cytoskeleton components, but some plasma membrane markers were detected. Though it would require genetic modification of the GBM cells, future studies could express an epitope tag on endolysosomes to facilitate rapid immunopurification prior to aptamer isolation.

Here we sought to address some of the questions raised by our previous selections (11, 12) and to test improved selection strategies toward functional aptamers and ApDCs that not only home to a tissue but deliver a relevant toxin cargo intracellularly. We first performed an unbiased cell-SELEX experiment with and without MMAE conjugation and demonstrated that *in vitro* selection could yield cell-specific aptamers regardless of toxin conjugation status. We showed that some aptamers bound target cells +/-MMAE regardless of whether they had originally been selected as toxin conjugates. However, we found that the ability to bind GBM cells *in vitro* after post-selection toxin conjugation did not translate into GBM tumor homing *in vivo* when aptamer biodistribution was measured 30 min post-i.p. injection. Unlike our prior results for *in vivo*-selected aptamers (11, 12) *in vivo GBM tumor homing of cell-selected aptamers was not observed regardless of conjugation status*. This result differs from reports for some cell- or protein-selected aptamers in other contexts (13, 14). We hypothesize that this difference results from the deliberately target-agnostic nature of our selections. In conventional SELEX, candidate aptamers are selected against specific internalizing receptors (e.g. EGFR or transferrin), known to be involved in BBB transcytosis. Although this approach has been effective, it does not enable the discovery of novel targets or novel aptamer delivery pathways, nor does it permit broad targeting across heterogenous tumor antigens. As such, it may deliver short-term efficacy, but its long-term potential is limited. We have therefore studied target-agnostic SELEX. We hypothesize that a weakness of target-agnostic cell- or *in vivo*-SELEX is that a majority of cell surface targets identified by aptamers do not result in internalization and lysosomal delivery required for toxin release by conventional mechanisms. Thus, there is impetus to adapt the unbiased selection approach to reward sub-cellular compartment-specific delivery. Our laboratory has previous investigated such strategies in other contexts (16–18).

Our new selections also explored the impact of timing on *in vivo* SELEX and whether selection could be improved by including rounds of *in vitro* cell-SELEX. We then devised a reward for intracellular delivery. Our results suggest that timing of aptamer of Ap-DC exposure *in vivo* is important. Increased circulation time had the apparent effect of reducing nonspecific background aptamer recovery, increasing selective pressure and thus the effectiveness of the selection. For the 30-min short-exposure library (Library #2), aptamer yield after lysosomal enrichment was comparable to bulk tumor enrichment. For this timepoint, four rounds of cell-SELEX after six rounds of *in vivo* SELEX were beneficial for improving *in vivo* homing. The 90-min exposure (Library #1) improved yield of aptamers recovered from the endolysosomal fraction and decreased background so improved enrichment was observed. Interestingly, for this timepoint, the addition of *in vitro* cell-SELEX rounds did not improve GBM tumor homing. However, it is important to acknowledge that these findings are limited to the rapid endolysosomal fractionation kit reward strategy employed in this *in vivo* SELEX.

It is also interesting to note that some top aptamers were observed in both the hybrid and *in vivo-*only selections, suggesting reproducibility of aptamer selection. In these cases, biodistribution patterns (i.e. bulk recovery from lung and kidney) differ for the same sequence in the final mouse of the respective selections, suggesting that the degree of tissue blood circulation may contribute to aptamer and Ap-DC biodistribution in the mouse. Unlike our previous *in vivo* DNA aptamer selections recovered four hours post-injection (background relatively low), shortening the circulation time to 30 or 90 min greatly increased the total undegraded aptamer dose still present in the mouse. Thus, we hypothesize that biodistribution pattern is dependent on circulation time.

There are many potential future directions for *in vivo* SELEX optimization. An intriguing finding from our work was the unusual occurrence of guanosine-rich aptamers for long-exposure Library #1. These sequences could be assayed for K^+^-dependent folding by electrophoretic mobility shift as evidence of G-quadruplex formation (19). Future studies should also evaluate the ApDCs selected in this selection for *in vitro* toxicity, binding, and internalization. Because the present work generated many anti-GBM aptamer and ApDC candidates, the field could benefit from determining if these strategies generated ApDCs with improved toxin delivery, homing, or target cell or tissue binding. It would also be pertinent to understand if hybrid selection involving *in vitro* and *in vivo* rounds gave rise to aptamers with improved tolerance to post-SELEX modification. Finally, *in vivo* selection for endolysosomal delivery should be repeated four hours post-injection to seek ApDCs with even improved tumor homing as had been observed in our original unconjugated DNA aptamer *in vivo* selections (11, 12)

## ACKNOWLEDGEMENTS

This work was supported by Mayo Clinic (L.J.M., J.N.S); L.J.M. is the Bernard Pollack Professor and J.N.S. is the William H. Donner Professor at Mayo Clinic. H3K27MPP cells were generously provided by Dr. Sameer Agnihotri at the University of Pittsburgh. Creation of the graphical abstract, Fig. 1AF, Fig. 2AB, Fig. 3A, and Supplementary Figs. S5 and S10 were facilitated by Biorender.com Doherty, C. (2026) https://BioRender.com/gu85a86.

## AUTHOR CONTRIBUTIONS

CDD, SJ, BAW, DMB, JNS and LJM conceived and designed the experiments. CDD, SJ, LLO, KKB, and BLC performed the experiments. CDD performed data analysis, CDD and LJM wrote the manuscript.

## SUPPLEMENTARY DATA

Supplementary Data are available with this deposit.

## DATA AVAILABILITY

All data are provided in supplementary materials. Constructs are available from the authors upon request.

