## Supplement for "Approaches to Optimize Cell Internalization and *In Vivo* Tumor Homing by Aptamer-drug Conjugates using SELEX"

### Table of Contents

**1 Supplementary methods**

**2 Supplementary data**

**Supplementary Table S1**. Experimental oligonucleotides for cell-SELEX.
**Supplementary Figure S1.** Aptamer strand purification by gel extraction during SELEX.
**Supplementary Figure S2.** Visualizing intracranial tumors with toluidine blue.
**Supplementary Figure S3.** Comparison of aptamer biodistribution +/- MMAE.
**Supplementary Figure S4**. Comparison of aptamer biodistribution +/- MMAE to *in vivo* selected ApDC candidates from previous work.
**Supplementary Figure S5.** Biodistribution of cell-selected aptamers +/-MMAE in mice bearing G43 flank tumors.
**Supplementary Figure S6.** Possible method to capture ApDCs that have undergone cleavage by cathepsin: use of 1-ethyl-3-(3-dimethylaminopropyl)carbodiimide) (EDC)- N-hydroxysuccinimide (NHS) coupling.
**Supplementary Figure S7.** Pilot studies attempting to capture 5'carboxyl aptamers with EDC-NHS coupling reaction.
**Supplementary Figure S8.** Conceptual method to capture cathepsin-cleaved aptamers during SELEX by size selection and gel purification.
**Supplementary Figure S9.** Endolysosomal isolation protocol to identify previous *in vivo*-selected ApDCs more likely to be internalizing.
**Supplementary Figure S10.** Lack of toxicity observed from former *in vivo*-selected ApDCs as determined by a CellTiter Glo assay 6 days post-treatment with ApDC, MMAE, or vehicle.
**Supplementary Figure S11.** Changes made for this in vivo SELEX compared to the first two done by our lab.
**Supplementary Figure S12.** Additional evaluation of the endolysosome extraction and libraries used during SELEX.
**Supplementary Figure S13.** Final nucleotide distributions in Library #1 (A, long exposure) or Library #2 (B, short exposure).
**Supplementary Figure S14.** Attempts to identify rare aptamers with improved tumor homing.
**Supplementary Figure S15.** Western blot of sample obtained from bulk tumor processing.
**Supplementary Figure S16.** Aptamer biodistributions after hybrid SELEX.

**3 Supplemental references**

### 1.0 Supplemental Methods

#### Random DNA library synthesis for cell SELEX

DNA oligonucleotides (**Supplementary Table S1**) were obtained from Integrated DNA Technologies. Two independent "orthogonal" libraries (library amplification requiring different 3′ DNA primers) carried 5′ modifications. One library was conjugated to MMAE (+MMAE library) and the other was conjugated to fluorescein (FAM; -MMAE library). Forward primers used during SELEX were synthesized and HPLC-purified either with 5′MMAE (IDT special order /5mod/; /5mod = NHS ester-PEG4-Val-Cit-PAB-MMAE (BroadPharm #: BP-25503) or 5′-fluorescein (/56-FAM) for +MMAE and -MMAE libraries, respectively. The reverse primer for the -MMAE library caried an internal 9-atom triethylene glycol spacer (/iSp9/) followed by a 3′-terminal 10-nt poly-A sequence that served as a non-extendable linker enabling electrophoretic separation of aptamers from complementary strands upon denaturing gel electrophoresis of the -MMAE library. For the +MMAE library, an unmodified reverse primer was sufficient because addition of 5′ MMAE created a distinguishable mobility shift in preparative polyacrylamide gels. Libraries were constructed with two constant 20-nt primer regions flanking 40 random nt. All oligonucleotides were reconstituted as 100 µM stocks. Primer stocks were 5 µM. To create both libraries for Round 1, the naïve library was amplified by five cycles of PCR using the following reagent volumes in a master mix: 320 µL 10× *Taq* polymerase buffer, 320 µL 1 mg/ml (10×) BSA, 256 µL 50 mM $MgCl_2$, 256 µL 2.5 mM dNTPs (dCTP, dATP, dGTP, and dTTP), 16 µL 100 µM library DNA, 320 µL 5 µM forward primer, 320 µL 5 µM reverse primer, 1347 µL water, and 45 µL *Taq* DNA polymerase (Invitrogen # 10342020). Reagents were mixed, aliquoted into 100 µL volumes in PCR tubes, and subjected to thermal cycling for five cycles using the following protocol: 95°C, 60 s; 6× (94°C, 15 s; 50°C, 35 s; and 72°C, 30 s); 4°C. Molar equivalent levels of primer to template were added in efforts to preserve library diversity and PCR cycles were limited to the minimum required to consume all forward primer with the goal of limiting PCR bias in the naïve library.

PCR reactions were pooled and precipitated from 2.5 volumes of ethanol after addition of glycogen carrier and sodium acetate to a final concentration of 0.3 M. After washing with 70% ethanol and air drying, PCR products were resuspended in water and deionized formamide (Ambion #AM9342), heated at 90°C for five min, and subjected to electrophoresis through a 7.5%

polyacrylamide gel [7.5 M urea, 19:1 acrylamide : bisacrylamide (Bio-Rad #1610144)] to separate the forward and reverse strands. For the +MMAE library, migration of the forward strand was retarded by the equivalent of ~10 nucleotides due to MMAE primer conjugation, permitting clear separation of the forward and reverse strands (**Supplementary Figure S1**). Forward strands of both libraries were visualized by UV shadowing (+MMAE at ~90 nt and -MMAE at 80 nt), excised with a razor blade, diced, and incubated in 0.3M NaOAc buffer at 56°C with agitation overnight. Eluted DNA was purified from gel fragments by passage through glass wool and precipitated from ethanol as described above. The concentration of the resulting naïve library was determined with a Nanodrop instrument at 260 nm using a library-specific estimated extinction coefficient (757,575 L/mol-cm). A 1-mL solution containing 800 pmol aptamer in PBS supplemented with 5 mM $MgCl_2$ and 4.5g/L glucose was created from the purified material. Libraries were heated at 90°C for 5 min and snap cooled on ice for 10 min. The 1 mL aptamer solution was added to a 1 mL solution of PBS containing 5 mM $MgCl_2$ plus 4.5 g/L glucose with 2× BSA and 2× tRNA (such that BSA and tRNA each had a final concentration of 0.1 mg/mL and the final aptamer concentration in the stock solution was 400 nM).

**Orthogonal Cell SELEX**

Libraries +/-MMAE were simultaneously subjected to the following SELEX protocol, although plates of cells and PCR reactions were manipulated separately. Prior to the day of selection, a 10-cm culture dish was seeded with either $4x10^6$ SVG-A cells or $1x10^6$ G43 cells. On the day of selection, SVG-A cells were first washed in wash buffer (PBS containing 5mM $MgCl_2$) and this wash buffer removed. The 2-mL solution of 400 nM aptamer library was added to negative selection (non-target) SVG-A cells with incubation at 37°C for 30 min, the dish being gently agitated every 10 min. The supernatant containing unbound aptamers was transferred from non-target SVG-A cells to washed and aspirated G43 cells (positive selection) with incubation at 37°C for 30 min, again with gentle agitation every 10 min. The supernatant containing unbound aptamers was aspirated from G43 cells and the dish washed 3× with wash buffer prior to cells being scraped in 1 mL wash buffer and collected. The dish was rinsed with an additional 400 µL wash buffer to collected any remaining cells. Collected cells in wash buffer were subjected to centrifugation at

500 × g for 5 min and the supernatant was aspirated. The resulting cell pellet was resuspended in 500 µL wash buffer and heated at 95°C for 10 min. The lysate was subjected to centrifugation at 13,100 × g for 5 min and the supernatant was recovered.

For Round 1, a full pool PCR was completed. To a 1-mL PCR reaction (100 µL 10× Taq pol buffer, 80 µL 2.5 mM dNTPs, 100 µL 10× BSA, 80 µL 50 mM MgCl2, 50 µL 10 µM forward primer, 50 µL 10 µM reverse primer, 20 µL water, and 20 µL Taq DNA polymerase), 500 µL of the recovered sample was added. The reaction was mixed and aliquoted into 10 individual 100-µL aliquots. Eight cycles of PCR were performed with the protocol above. Reactions were pooled and 210 µL of the reaction was used as template in the following PCR.

For all subsequent rounds, 210 µL of lysate was used in 2.1 mL PCR reactions (210 µL 10× Taq pol buffer, 168 µL 2.5 mM dNTPs, 210 µL 10× BSA, 168 µL 50 mM $MgCl_2$, 210 µL 5 µM forward primer, 210 µL 5 µM reverse primer, 672 µL water, and 29.4 µL Taq). A pilot PCR reaction was performed with 100 µL of the aliquot from the SELEX PCR reaction and a separate 100 µL water background reaction (10 µL 10× Taq pol buffer, 8 µL 2.5 mM dNTPs, 10 µL 10× BSA, 8 µL 50 mM $MgCl_2$, 10 µL 5 µM forward primer, 10 µL 5 µM reverse primer, 42.6 µL water, and 1.4 µL Taq). The 100-µL reactions were divided into five 20-µL reactions and subjected to the following protocol: 95°C, 60 s; 6× (94°C, 15 s; 50°C, 35 s; and 72°C, 30 s); 4°C. Aliquots were removed at every few cycles (i.e. 0, 14, 18, 22, 26 cycles) and 2 µL of each reaction was mixed with 10 µL formamide (Ambion, #AM9342), heated at 90°C for 5 min, 2 µL of loading dye added, and samples were subjected to electrophoresis through 7.5% denaturing polyacrylamide gels [7.5 M urea, 19:1 acrylamide : bisacrylamide (Bio-Rad #1610144)]. Gels were stained with SYBR gold (Invitrogen, #S11494) and imaged. The optimal PCR cycle was selected based on the disappearance of the primers and the clear appearance of the desired amplicon with as little non-specific products and water background signal as possible. Aliquots of the main preparative SELEX PCR were then subjected to amplification for this optimal number of cycles. Reactions were then pooled and precipitated from ethanol as described above. Amplicons were resuspended in water and formamide and heated at 90°C for 5 min. The aptamer strand was isolated by gel purification through a 7.5% polyacrylamide gel and eluted in 0.3M NaOAc overnight at 56°C with agitation. Apatmer DNA was purified from the gel fragments as described above. Aptamers were resuspended in water and the resulting concentrations for each library calculated from the absorbance at 260 nm as determined by a nanodrop instrument using estimated extinction coefficients. For subsequent rounds of selection 100 pmol of each library was heated and snap-cooled in 1 mL PBS containing 5 mM $MgCl_2$ and 4.5 g/L glucose. This 1 mL was added to another 1 mL of PBS containing 5 mM $MgCl_2$ and 4.5 g/L glucose containing 2× competitors (tRNA and BSA) so the final tRNA and BSA concentrations were 0.1 mg/mL.

**Mice**

Female athymic nude (Strain #553) mice were purchased from Charles River Laboratories. All animal experiments were approved by Mayo Clinic Institutional Animal Care and Use Committee Protocol #A00006585-22. Orthotopic tumor inoculation and bioluminescence imaging were performed as previously described (1, 2).

**In vivo SELEX in orthotopic PDX mice**

Prior to SELEX, two cohorts of mice were injected with explants of GBM43-eGFP-FLUC2 cells. In the first two cohorts, 12 mice were injected in three groups (four mice per group) with the following cell counts: 100,000 cells, 25,000 cells, or 10,000 cells. In the third cohort, 6 mice were injected in four groups (two mice per group) with the following counts: 100,000 cells, 10,000 cells, or 1,000 cells. This procedure distributes tumor development over time. Tumor development was monitored by bioluminescence imaging (BLI) (IVIS Spectrum). Once the tumor reached a BLI reading between $5\times10^8$ and $1\times10^9$, the mouse was enrolled for a round of *in vivo* SELEX.

On the day of selection, one mouse received an intraperitoneal (i.p.) injection of 250 pmol long-exposure aptamer library (Library #1) in 250 µL PBS containing 1 mM $MgCl_2$ 90 min prior to harvest, followed by a second i.p. injection of 250 pmol of short-exposure aptamer library (Library #2) 30 min prior to harvest. At harvest, the mouse was euthanized with $CO_2$ and subjected to scrupulous transcardial perfusion with 30-60 mL PBS as follows: the euthanized mouse was pinned outstretched at a vertical angle and a 21g × 1″ needle was inserted into the left ventricle while the heart was still beating. The mouse was perfused until the liver changed from dark red to pale brown with the effluent gravitationally pooling below the mouse. The brain was extracted, and the $GFP^+$ tumor was dissected with the aid of fluorescent goggles. Extreme care was used to avoid aptamer contamination of surfaces and instruments.

Only at the final round were all organs collected for aptamer extraction and deep sequencing. The brain tumor was divided in half: half was immediately cooled on wet ice and processed normally as described below and the second half was snap frozen on dry ice and saved for bulk tissue processing with other organs (**Figure 3A**). The adjacent brain, cerebellum, heart, lungs, liver, kidneys, spinal cord, and sciatic nerve were also extracted and snap frozen. The bone marrow was collected from both femurs by centrifugation at $\geq$10,000 × g. All tissues were snap-frozen on dry ice and stored at -80$^{o}$ C.

**Rewarding aptamer internalization**

For selections to reward aptamer internalization, tissue was subjected to an endolysosomal compartment enrichment protocol using a Lysosomal Isolation Kit (Abcam, #ab234047). After tumor harvest, the fresh tissue was immediately placed in a tared tube containing 500 µL Abcam Lysosome Isolation Buffer and kept on wet ice. The tissue was weighed and the Lysosome Isolation Buffer volume increased accordingly to the optimal ratio according to the manufacturer's instructions. Protease Inhibitor Cocktail was added, and the tissue was gently mechanically dissociated using a plastic pestle and the resulting slurry transferred by pipet to a Dounce homogenizer and subjected to 12 strokes. The homogenate was subsequently processed according to the manufacturer's protocol. All steps were done on ice in a cold room at 4$^{o}$C. Following the PBS wash and centrifugation in 18,000 × g, the endosomal fraction was resuspended in 60 µL (75 µL for Round 1) of PBS. A 30 µL aliquot (60 µL for Round 1) was mixed with an equal volume of lysis buffer (300 µL: 50 mM Tris HCl, pH 7.4, 150 mM NaCl, 1% NP-40, 0.5% Na Deoxycholate, 1 mM EGTA, and 1 mM NaF) and mixed by vortex. The lysate was heated at 95$^{o}$C for 10 min and subjected to centrifugation at 13,100 × g for 5 min.

For Round 1, the library was pre-amplified. A 1 mL PCR reaction was created (100 µL of 10× Taq polymerase buffer, 80µL of 2.5 mM dNTPs, 100 µL 10× BSA, 80 µL 50 mM $MgCl_2$, 50 µL each of both forward primers, and 50 µL each of both reverse primers, 365.9 µL water, 60 µL of template, and 14.1 µL of Taq (Invitrogen #10342020).). Six cycles of PCR were performed. To the 3.2 mL PCR reactions (one for each library) below, 320 µL of the PCR reaction was added.

For all subsequent rounds, a 30-µL sample of the crude tumor endolysosome-enriched lysate was added to each of the two 3.2-mL PCR reactions (one for each library) comprising 320 µL 10× Taq polymerase buffer, 320 µL 10× BSA, 256 µL 50 mM $MgCl_2$ (Library #1 64 µL 50 mM $MgCl_2$ and 192 µL of water), 256 µL 2.5 mM dNTPs (2.5 mM dCTP, dATP, dGTP, and dTTP), 320 µL 5 µM forward primer, 320 µL 5 µM reverse primer, 1333 µL water, and 45 µL *Taq* DNA polymerase (Invitrogen #10342020). Note that for this selection the reverse primer was alternated every round between three primers [5′ FAM 20-nt unmodified primer, 5′FAM primer with a non-extendable linker and 3′-terminal $(dA)_{30}$ sequence (50 nt total), and a 5′FAM-modified primer with a non-extendable linker followed by a 3′-terminal $(dA)_{36}$ sequence (56 nt total)] (**Supplementary Figure S1B**). Toggling avoided parasitic PCR products. Cycle optimization was performed as above and samples obtained at rounds 0, 25, 30, 35, and 40. The 100-µL reactions were divided into five 20 µL reactions and subjected to the following protocol: Library #1: 95°C, 60 s; 6× (94°C, 15 s; 54°C, 35 s; and 72°C, 30 s); 4°C and Library #2: 95°C, 60 s; 6× (94°C, 15 s; 51°C, 35 s; and 72°C, 30 s); 4°C. Library #1 became increasingly more difficult to amplify over selection rounds, which necessitated an increase in the annealing temperature, decrease in the concentration of $MgCl_2$, and/or reliance on FailSafe Buffer PCR optimization (Biosearch Technologies, #FS9901K). The remaining volume was subjected to PCR amplification at the optimal number of cycles determined for that round (usually ~30-35 cycles). PCR reactions were pooled, precipitated from ethanol after addition of NaOAc and glycogen as before, and resuspended in water and deionized formamide (Ambion #AM9342). Samples were heated at 90°C for 5 min prior to loading on a pre-run 7.5% denaturing polyacrylamide gel [7.5 M urea, 19:1 acrylamide : bisacrylamide (Bio-Rad #1610144)] and subjected to electrophoresis at 600 V for 2.5 h. The forward non-fluorescein-labeled strand was visualized by shadowing with a UV lamp over a fluorescent TLC plate and excised with a fresh razor blade (**Supplemental Figure S1B**). This band was diced and transferred to microcentrifuge tubes with 0.3M NaOAc buffer and incubated overnight at 56°C with agitation at 1500 rpm. Polyacrylamide fragments were removed as before. ApDCs were extracted from the solution by precipitation from ethanol as before and resuspended in 80-100 µL water. The absorbance at 260 nm (Nanodrop) was divided by the molar extinction coefficient to calculate the concentration of the recovered library. The sample for the next round of selection was prepared by adding 28 µL 10× PBS, 2.8 µl, 280 pmol of library, and increasing the volume to 280 µL with water. This cycle was repeated for each round of selection.

**Hybrid SELEX**

After 6 rounds of *in vivo* SELEX performed as described above, libraries were divided: half continued to 10-11 rounds of *in vivo* SELEX only and half underwent 4 rounds of cell SELEX followed by one final SELEX round *in vivo*.

The four cell-SELEX rounds were performed as follows. One T175 flask was seeded with $10^7$ cells and allowed to grow until 90-95% confluent (~2 $\times 10^7$ cells). On the day of selection, 5 mL of selection buffer (PBS containing 1 mM $MgCl_2$ and 4.5 g/L glucose) was supplemented with a final concentration of 0.1 mg/mL sheared salmon sperm DNA (produced by three cycles of 2 min sonication at power setting 10; Sonic Dismembrator 60, Fisher Scientific) and 0.1 mg/mL BSA. Cells were washed once with selection buffer and the selection buffer containing sheared salmon sperm DNA and BSA was added, to which 100 µL of snap-cooled Library #1 in PBS containing 1 mM $MgCl_2$ was added. Cells were incubated at 37°C for 30 min and then 100 µL of snap-cooled Library #2 in PBS containing 1 mM $MgCl_2$ and was returned to the incubator for 15 additional min (total incubation times: Library #1: 45 min; Library #2: 15 min). Cells were washed 3 times in selection buffer and then briefly trypsinized followed by the addition of 10 mL of media. Cells were collected in a 50 mL conical tube and subjected to centrifugation at 600 × g for 10 min according the Abcam Lysosomal Isolation kit protocol. The manufacturer's protocol for cells was followed for remaining steps. Cycle optimization was performed as described above, except that cycles 0, 21, 24, 27, 30, were evaluated. The remaining steps of selection were performed as described above except that 100 pmol of ApDC library was prepared and used for each round for cell SELEX. For the final *in vivo* round of the hybrid selection, the above *in vivo* SELEX protocol was performed and 280 pmol/280 µL were used for i.p. injection.

**Confirmation of library conjugation**

To confirm prepared libraries were successfully conjugated to MMAE, 1pmol ApDC library was incubated with or without anti-MMAE antibody (Creative Diagnostics # DMABA-JX124) at 1:10 dilution for 1 h at room temperature. Solutions were subjected to electrophoresis through 8% native polyacrylamide gels (29:1 acrylamide:bisacrylamide Bio-Rad #1610146) and stained with SYBR gold (Invitrogen #S11494).

**Confirmation of library size**

To confirm the size of the DNA library, 1 pmol of library was subjected to electrophoresis through a 7.5% denaturing polyacrylamide gel [7.5 M urea, 19:1 acrylamide : bisacrylamide (Bio-Rad #1610144)] and/or 1 pmol of duplex was subjected to electrophoresis through an 8% native polyacrylamide gel (29:1 acrylamide:bisacrylamide Bio-Rad #1610146). The gel was stained with SYBR gold (Invitrogen #S11494) and imaged.

**Aptamer sample preparation for deep sequencing**

Final library rounds for both the *in vivo* only selection and the hybrid selection (Round 10C and Round 11C) were sequenced initially for rapid results using AmpliconEZ Seq through Azenta. Libraries from the final round were subjected to 6 cycles of PCR in two 100-µL reactions per library. The PCR product was purified with a Qiagen MinElute Purification Kit (Qiagen #28004) and eluted in 20 µL water. DNA concentration was measured by Qubit using 3 µL sample. Duplex size and quality were confirmed by electrophoresis through an 8% native polyacrylamide gel (29:1 acrylamide:bisacrylamide; Bio-Rad #1610146). Samples (20 ng/µL in 25 µL) were then submitted to the vendor for sequencing.

To appreciate the biodistribution of top sequences, only *in vivo* SELEX libraries Rounds 5, 7, 9, and the final round were submitted for deeper Illumina sequencing in the Mayo Clinic Genomics Core by MiSEQ technology. Libraries obtained from these rounds were subjected to 6 rounds of PCR to create duplexes. Libraries #1 and #2 for a given round were combined at this step and were purified with the Qiagen MinElute Purification Kit (Qiagen #28004) and eluted in 20 µL water. Concentration was measured with Qubit.

To further determine the biodistribution of the sequences in the library, aptamers were extracted from the final SELEX rounds of both the *in vivo*-only SELEX library mouse (Round 10/11) and the hybrid SELEX library mouse (Round 11C). To increase the yield of aptamers from these bulk frozen tissues, an optimized aptamer extraction method was employed (3). Bone marrow samples were rinsed with 500 µL PBS and subjected to centrifugation at 200 × g for 5 min to remove lysed blood cells and then processed as for other tissue samples. Samples of mass less than 200 mg were homogenized in 300 µL Qiagen Plasmid Extraction Kit Buffer P1 with a plastic pestle. Lysis buffer (300 µL: 50 mM Tris HCl, pH 7.4, 150 mM NaCl, 1% NP-40, 0.5% Na Deoxycholate, 1 mM EGTA, and 1 mM NaF) was then added with incubation on wet ice for 3 h. Samples were sonicated at power 8 (Sonic Dismembrator 60, Fisher Scientific) for 10 s three times with 30-s of cooling on ice between treatments. Samples were heated to 95$^{o}$ C for 10 min and subjected to centrifugation at 13,100 × g for 5 min. The supernatant was removed and analyzed by qPCR as described above.

Libraries were amplified for the optimal number of cycles determined by analytical PCR in each selection round using unmodified primers. Duplexes were created from the libraries with PCR reactions containing: 10 µL 10× *Taq* DNA polymerase buffer, 10 µL 10× 1 mg/mL BSA, 8 µL 50 mm $MgCl_2$, 8 µL 2.5 mm dNTPs, 10 µL library, 10 µL 5 µM forward primer, 10 µL 5 µM reverse primer, 32.6 µL water, and 1.4 µL *Taq* DNA polymerase. Six rounds of PCR were performed and then 20 µL fresh master mix was added to reach sample for one additional cycle to ensure fully-extended duplexes (4) PCR product size and quality were confirmed by electrophoresis through an 8% native polyacrylamide gel (29:1 acrylamide:bisacrylamide; Bio-Rad #1610146). Respective bar-coded libraries derived from the same organ and mouse were then combined.

PCR products were purified with the Qiagen MinElute Purification Kit (Qiagen #28004). Ten nanograms of eluted DNA, as quantified by the Qubit HS Duplex DNA Quantification Kit (Invitrogen #Q32851), were used as input for the NEBNext Ultra II DNA Library Prep Kit (NEB #E7645S) using associated NEBNext Multiplex primers index (primers set 1 and set 2; NEB, E7335S and E7500S). After preparing samples according to the manufacturer's instructions except that recommended volumes were halved and in the purification of adaptor ligated DNA Qiagen MinElute Purification kit was used instead of beads. Samples were then assessed by gel electrophoresis through an 8% native polyacrylamide get (29:1 acrylamide:bisacrylamide) prior to analysis by high-throughput paired-end 150 cycle sequencing on a single lane of an Illumina MiSeq instrument (Mayo Clinic Sequencing Core).

**Sequence Analysis**

Sequence files from Azenta were received in .gz format. Sequence files from the Mayo Clinic Sequencing Core were obtained in .cram format and converted to .bam format and then to .fastq format. Usearch was used to first merge paired-end reads. Merge files were filtered using the usearch maxee function, discarding files with errors >0.5. Reverse strands were isolated, and reverse complements created using the SeqKit reverse complement tool. The two files were combined with the simple concatenate function in Linux. AptaSUITE was used to filter any reads that did not contain both forward and reverse primers within an error of 3 bases, and aptamers ranked by sequence or cluster abundance (5).

Following deep sequencing analysis, aptamer enrichment was analyzed by identifying the most prevalent sequences and sequence clusters at Round 10 and by determining area under the curve for all sequences identified at Round 10. Individual candidate aptamers were then synthesized by IDT and converted to 5′MMAE ApDCs by in-house PCR and gel purification. Biodistribution of individual ApDCs was estimated by equation (1):

$$Molecules\ of\ aptamer_{i,tissue} = fraction\ of\ pool_i \times \frac{total\ molecules_{tissue}}{mass_{tissue}} \quad (1)$$

where $i$ indicates a given individual aptamer, and *fraction of the pool*$_i$ is calculated by number of counts for aptamer $i$ divided by the total of number of counts in the pool. The total number of molecules was determined by qPCR analysis of the library in every tissue sample. Recovered fraction of in tumor relative to all organs at time of harvest for aptamers was calculated by equation (2):

$$Fraction\ of\ injected\ dose_{i,tissue} = \frac{Number\ of\ molecules_{i,tissue}}{\sum_j Number\ of\ molecules_{i,j}} \quad (2)$$

### 2.0 Supplemental Data

**Supplemental Table S1. Experimental oligonucleotides for cell-SELEX**

| Serial number | Description | Sequence |
|---|---|---|
| LJM-6828 | +MMAE cell-SELEX library | AGTCTGTTCTCCTGTCTCAG ($N_{40}$) TGAGGTTACCTGGTGAACTG |
| LJM-7166 | +MMAE forward SELEX primer | /5VCPMPEG4N/AGTCTGTTCTCCTGTCTCAG |
| LJM-6882 | +MMAE reverse SELEX primer | CTACCACAGTCCCAAGATGT |
| LJM-7167 | -MMAE cell-SELEX library | /56-FAM/ATAGATGCGGAGATTATGGC ($N_{40}$) GTGTGACCCGTTATGCTCGA |
| LJM-7168 | -MMAE forward SELEX primer | /56-FAM/ATAGATGCGGAGATTATGGC |
| LJM-7169 | -MMAE reverse SELEX primer | AAAAAAAAAAAAAAAAAAAA/iSp9//iSp9/TCGAGCATAACGGGTCACAC |
| **7449** (ApDC 1) | Selected with conjugation (MMAE-6828 library) | AGTCTGTTCTCCTGTCTCAGGACCAACGCTTGAGTATGGACAGACGTGACCGTCTCGGCGACATCTTG GGACTGTGGTAG |
| **7450** (ApDC 2) | | AGTCTGTTCTCCTGTCTCAGTAAACTGGGCCGTAGGGACCATCTTACCTGTTGCGGATCAACATCTTG GGACTGTGGTAG |
| **7451** (ApDC 3) | | AGTCTGTTCTCCTGTCTCAGGCCGGAGGGACTGAAATGCGACGTTTACCGTGCGTAGACCACATCTTG GGACTGTGGTAG |
| **7546** | Neg Control (MMAE-6828 library) | AGTCTGTTCTCCTGTCTCAGGGATCTTAGCGGTTTAGCGGATGATTATCGAGACCGCGTCACATCTTG GGACTGTGGTAG |
| **7452** (Ap 1) | Selected without conjugation (LJM-7167 library) | ATAGATGCGGAGATTATGGCGGACTGCAGAGACCACCTGACGTTGCGCATTGTGACCGGTGTGTGACC CGTTATGCTCGA |
| **7453** (Ap 2) | | ATAGATGCGGAGATTATGGCATGTGGACTGCAGAGACCGCTGGCTAAGTGTGAGTTGCTGGTGTGACC CGTTATGCTCGA |
| **7454** (Ap 3) | | ATAGATGCGGAGATTATGGCAGGCCGCAGGGATCTACCTTTCTACGTATCCGAAAGATGTGTGTGACC CGTTATGCTCGA |
| 7547 | Neg Control (LJM-7167 library) | ATAGATGCGGAGATTATGGCGTTGGTACTTGTGAGTCTCAGGCCAGCCCCATGCGAACACGTGTGACC CGTTATGCTCGA |

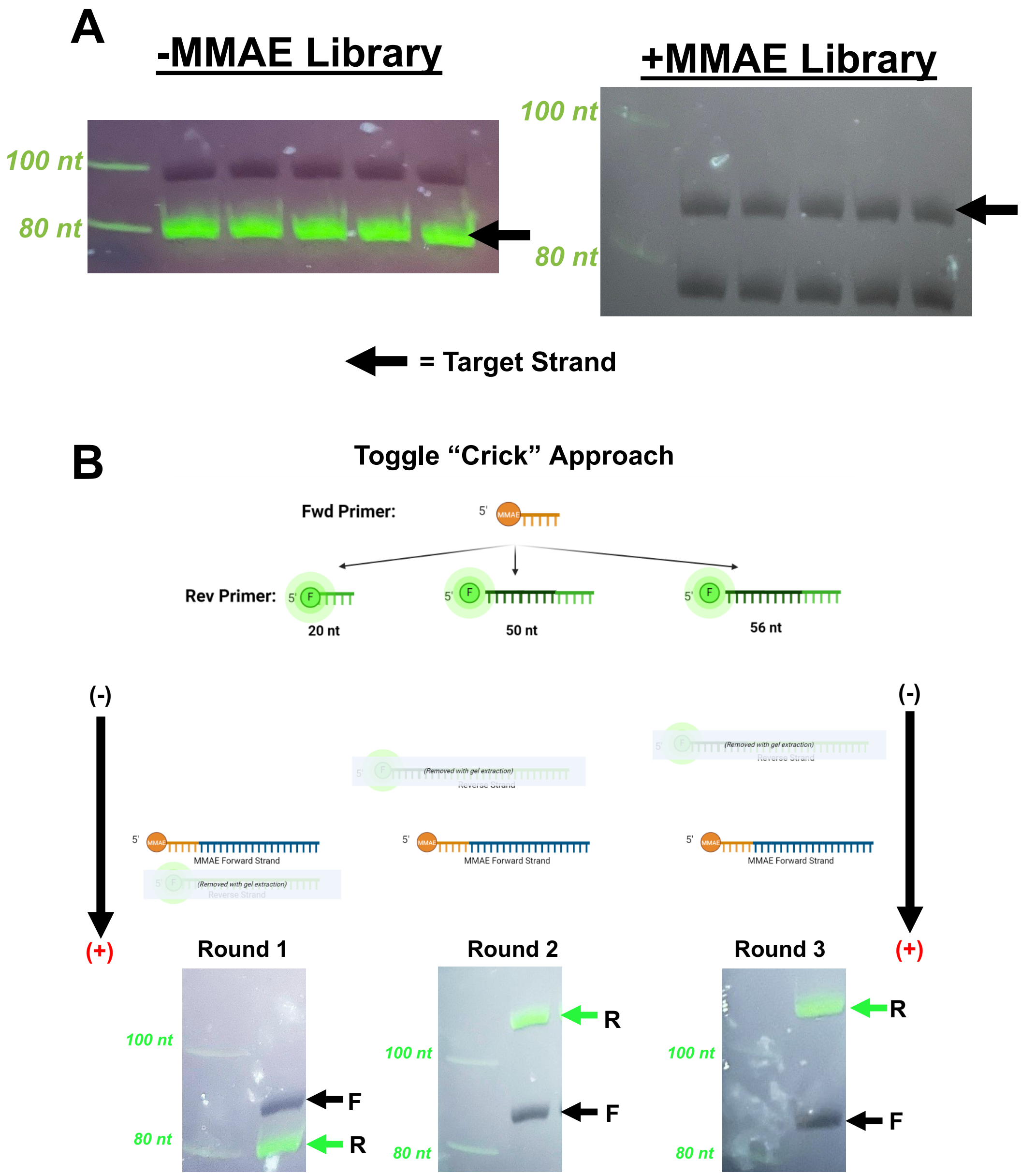


**Supplementary Figure S1. Aptamer strand purification by gel extraction during SELEX.** The forward (aptamer) strand of the PCR duplex is purified by denaturing polyacrylamide gel electrophoresis and gel extraction. A. For cell-SELEX, the -MMAE library utilized a 5'FAM label on the forward strand. The reverse strand was synthesized with a 3' $dA_{20}$ sequence downstream of a non-extendable linker. Isolation of MMAE-conjugated library sequences relied on the mobility retardation due to MMAE conjugation. B-C. For *in vivo* SELEX, a "toggle Crick" protocol was developed to assure proper strand identification after PCR. The reverse strand was labeled with 5'FAM with PCR alternating between three primers of different sizes: 20 nt (no non-extendable linker), 50 nt (non-extendable linker upstream of $(dA)_{30}$ terminus) or 56 nt (non-extendable linker upstream of a $(dA)_{36}$ terminus).

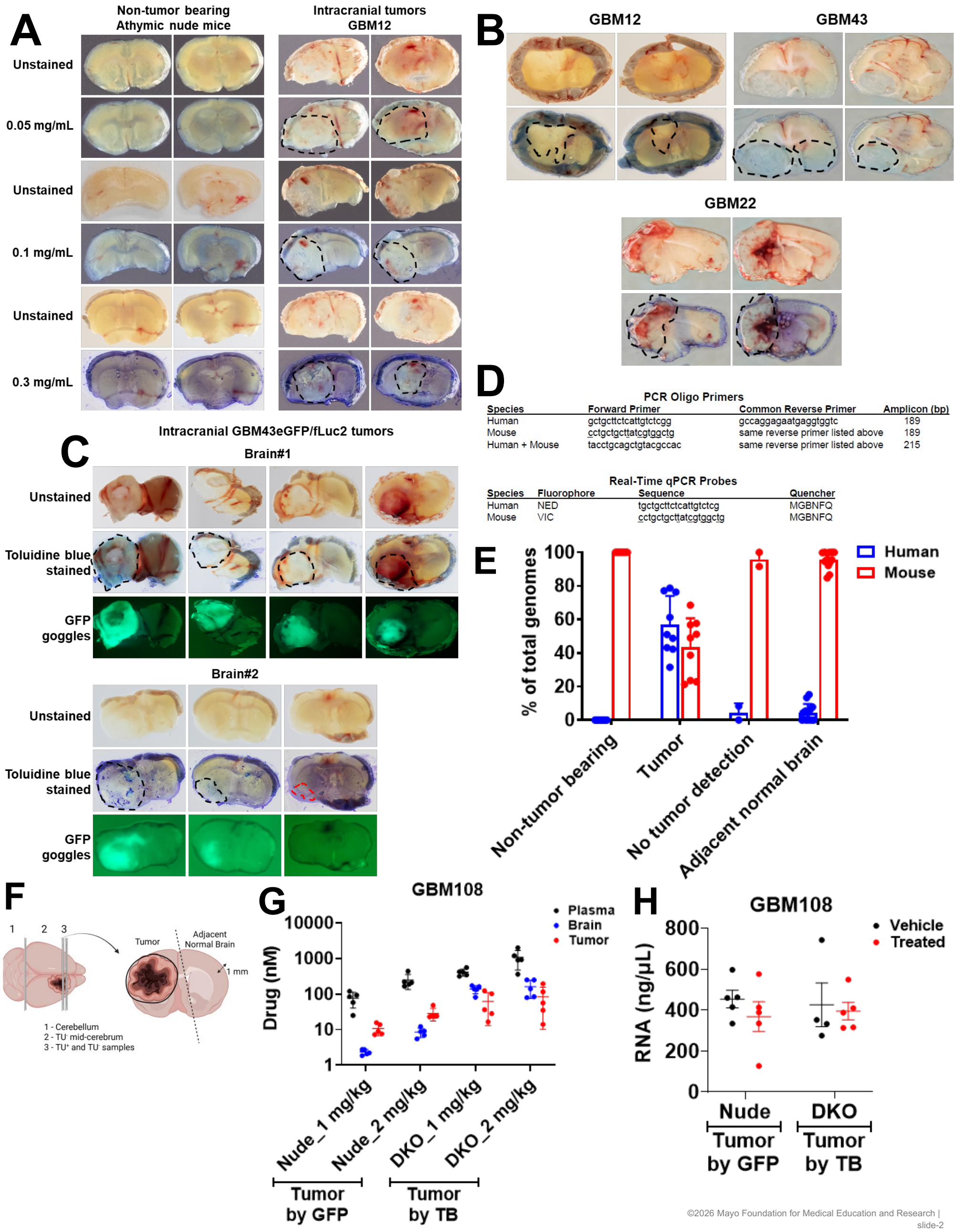

©2026 Mayo Foundation for Medical Education and Research | slide-2

**Supplementary Figure S2. Visualizing intracranial tumors with toluidine blue (TB).** Intracranial tumors distinguished from surrounding normal brain after staining with TB, an acidophilus metachromatic nuclear stain used in histology and vital staining. TB has been used to detect mucosal abnormalities and identify carcinoma of the oral cavity. A. TB detection of GBM12 tumors under different concentrations. B. TB staining aids tumor visualization in brain slices from several Mayo GBM PDXs. C. GBM43eGFP/fLuc2 intracranial tumor area delineated by TB staining matched with the tumor area detected by GFP fluorescence using GFP illumination. (D-F). Tumor and normal brain samples from diverse PDX models, collected after TB staining, showed abundant presence of human and mouse PTGER2 genes, respectively, as determined by PCR analysis of genomic DNA. D. Primers were used from Alcoser et al. (6). E. qPCR results for human vs mouse. Each dot represents a sample from an individual mouse. F. Processing of GBM tumors harvested after TB staining for various downstream analyses. (G-H). Using LC-MS/MS, concentration of drug was measured in the plasma, normal brain and tumor samples (TB or GFP guided) collected after brigimadlin treatment of nude mice with GBM108 parental tumor or double knockout (DKO) mice lacking efflux pumps with intracranial GBM108eGFP/fLuc2 tumor. F. Using nanodrop, concentration of RNA was determined in tumor samples collected by GFP signal or by TB staining from the untreated or brigimadlin-treated GBM108eGFP/fLuc2 or GBM108 parental tumors, respectively. Each dot represents data from an individual mouse.

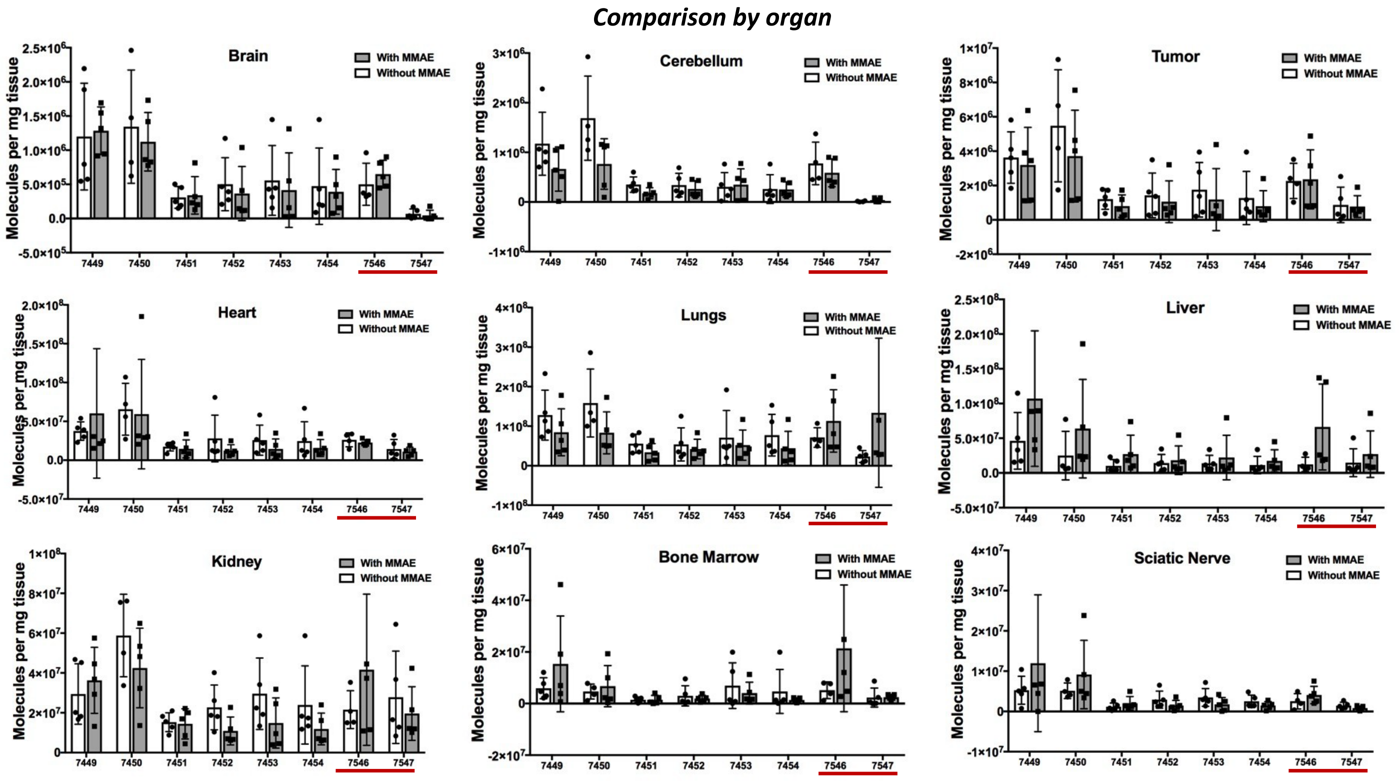


**Supplementary Figure S3. Comparison of aptamer biodistribution +/- MMAE.** Data are as shown in Figure 1H-I, but depicted per sequence per organ to highlight any difference between aptamers +/-MMAE. Negative control sequences are 7546 (scramble from the +MMAE selection) and 7547 (scramble from the -MMAE selection).

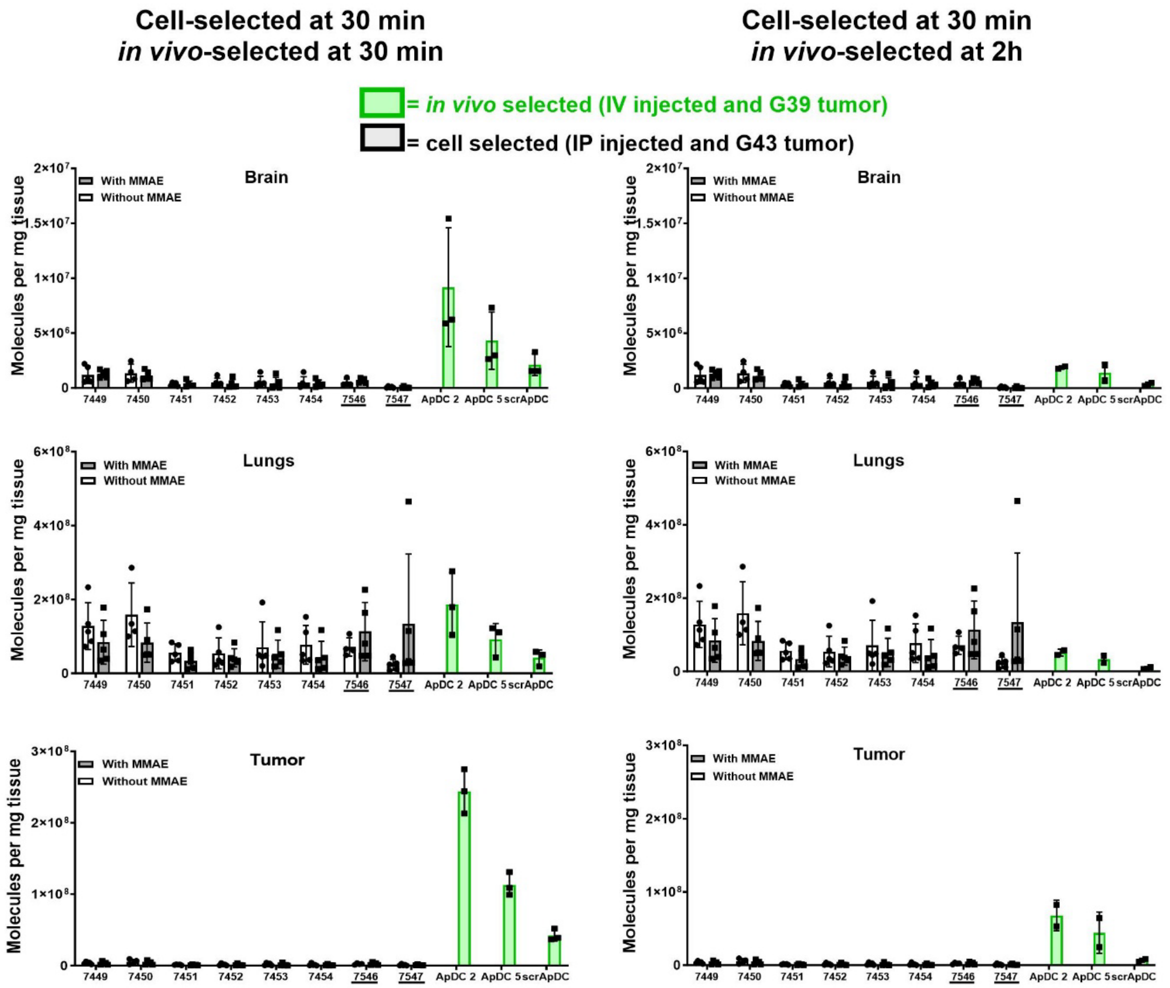


**Supplementary Figure S4. Comparison of +/- MMAE aptamer biodistribution in brain, lungs, and tumor to *in vivo*-selected ApDC candidates from previous work Doherty et al., 2026** (7). Same data as shown in Figure 1H-I, but graphed per sequence per organ of specific interest (brain, lung, and tumor) to demonstrate the lack of any difference between +/-MMAE versions of sequences. Results from the biodistribution at 30 min of *in vivo* selected ApDCs in Doherty et al. 2026 (green) are added for comparison. Note that the *in vivo* selected ApDCs were handled differently in that they were (1) intravenous (i.v.) injected into mice bearing (2) a G39 tumor, as they were selected on G39 tumors. Given the comparison between intraperitoneal (i.p.) and i.v. injections done in Doherty et al., 2026, there is significant evidence that the *in vivo* selected candidates have similar biodistributions to the tumor whether they are i.p. or i.v. injected (7). Negative control sequences are 7546 (scramble from the +MMAE selection) and 7547 (scramble from the -MMAE selection).

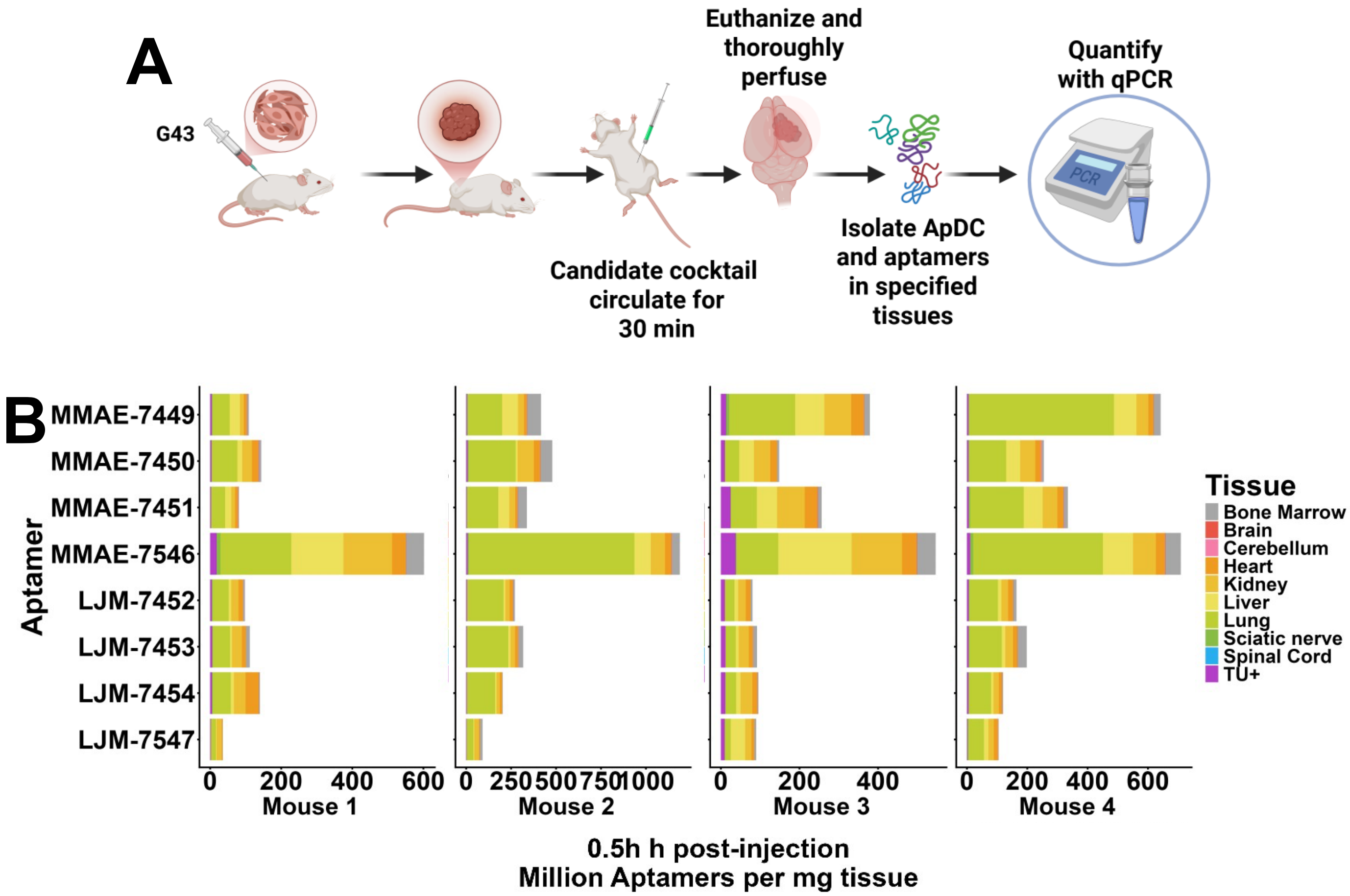


**Supplementary Figure S5. Biodistribution of cell-selected aptamers +/-MMAE in mice bearing G43 flank tumors.** This study was identical to the study shown in Figure 1H-I, except mice bore flank tumors to understand if poor biodistribution reflects the BBB or poor tissue homing *in vivo*. A. Schematic diagram of the experiment evaluating cell-selected aptamer or ApDC candidate biodistribution in mice bearing flank PDX GBM tumors B. Cell-SELEX derived aptamers were evaluated for their respective biodistributions with and without MMAE in mice with flank tumors. Two cocktails were created: Cocktail 1 contained the selected versions of the candidates and Cocktail 2 contained these candidates in the opposite MMAE conjugation state Mice received an I.P. injection of 400 µL (50 pmol/candidate) of either cocktail. After 30 min post-injection, the mice were euthanized and thoroughly perfused. Individual aptamers were quantified in the tissues using sequence-specific PCR primers (B).

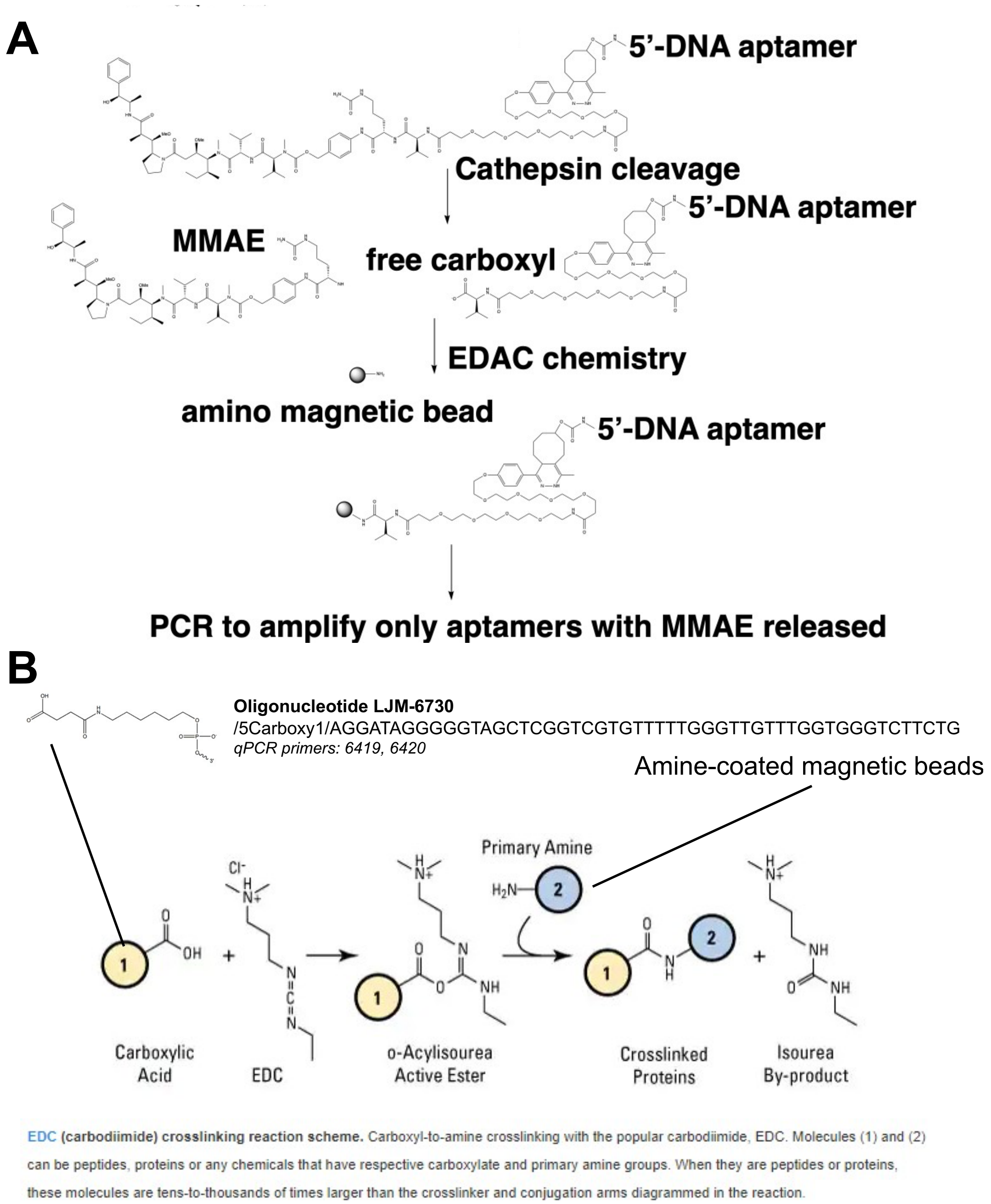


**Supplementary Figure S6. Possible method to capture ApDCs that have undergone cleavage by cathepsin: use of 1-ethyl-3-(3-dimethylaminopropyl)carbodiimide) (EDC)- N-hydroxysuccinimide (NHS) coupling.** A. Schematic diagram of concept. Following cathepsin cleavage, the aptamer will possess a unique 5'carboxyl group, amenable to reaction with EDC-NHS to conjugate the aptamer to an amino-modified magnetic bead to allow selective PCR after stringent washing. B. Diagram adapted from Thermo Fischer website (https://www.thermofisher.com/us/en/home/life-science/protein-biology/protein-biology-learning-center/protein-biology-resource-library/pierce-protein-methods/carbodiimide-crosslinker-chemistry.html) showing the intended reaction.

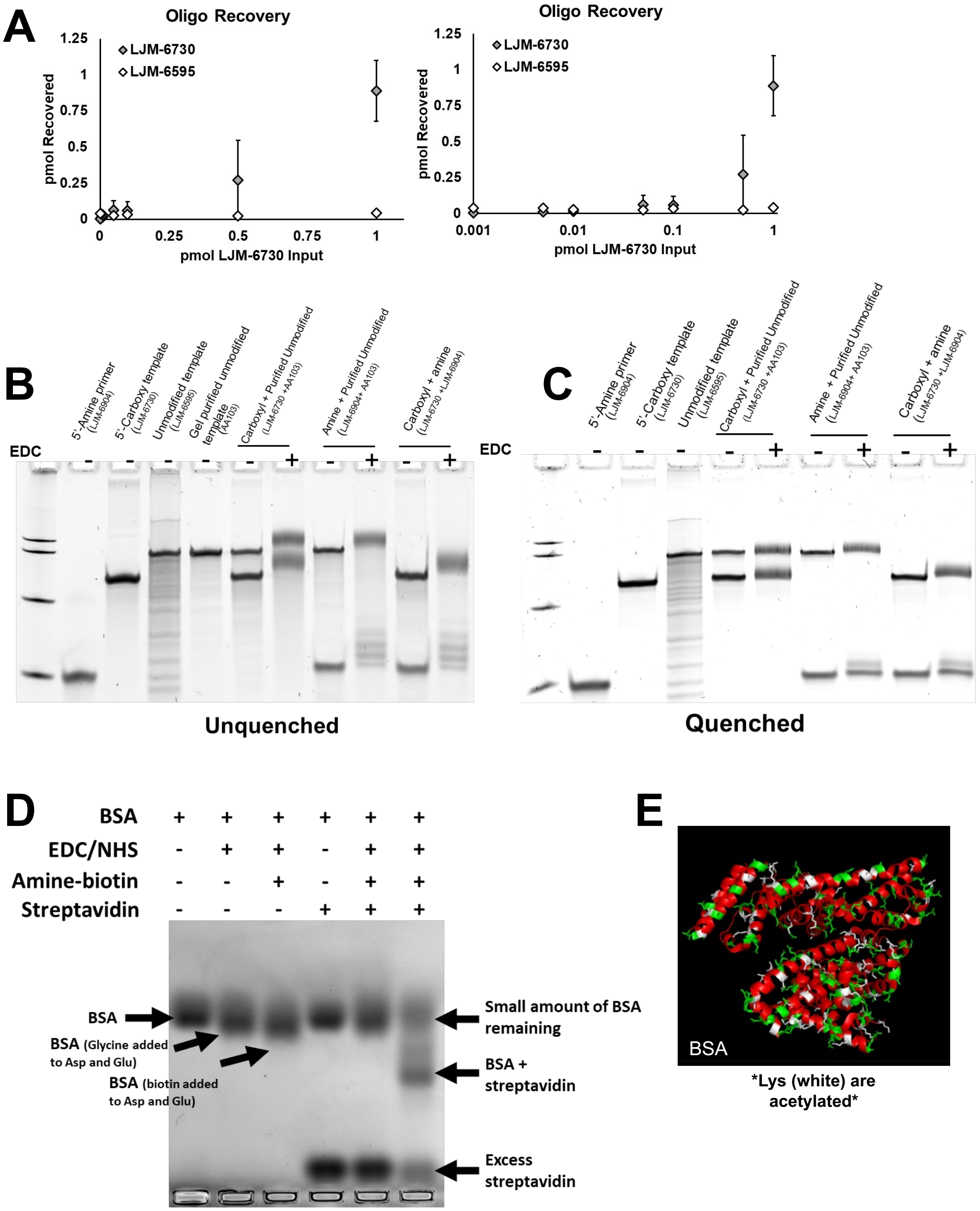


**Supplementary Figure S7. Pilot studies attempting to capture 5'carboxyl aptamers with EDC-NHS coupling reaction.** A. Replicate studies capturing a 5'carboxyl aptamer (LJM-6730) in the presence of a 5'hydroxyl aptamer (LJM-6595) as detected by qPCR. The reaction is successful for higher concentrations of 5'carboxyl aptamer, but yield is poor at lower concentrations necessary for SELEX. (B-C). A simplified reaction was attempted to monitor efficiency by coupling a 5'amino primer (20 nt) to a 5'carboxyl template. The reaction was completed with (B) and without (C) quenching. No evidence of coupling was observed. (D-E) As EDC-NHS coupling reactions are commonly used in protein conjugation, we sought to validate the efficacy of the reaction by coupling bovine serum albumin (BSA) lysine residues to an aminobiotin conjugate and identifying products using a agarose gel mobility shift assay in the presence of streptavidin. Despite numerous surface lysine residues, only ~ 60% of BSA was conjugated to biotin under these conditions, an efficiency insufficient for the intended SELEX capture application.

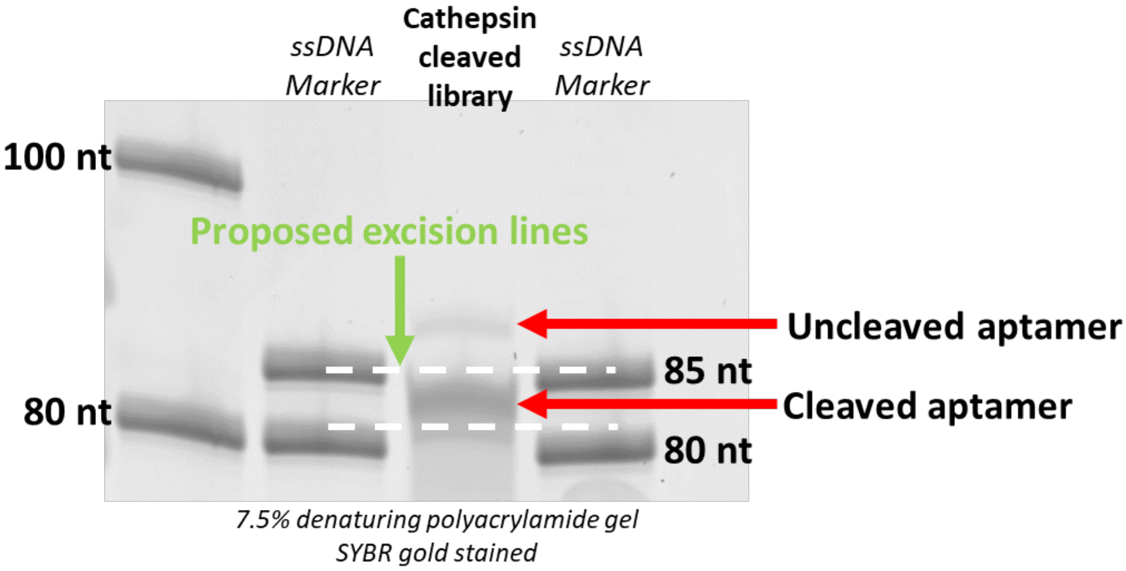


**Supplementary Figure S8. Conceptual method to capture cathepsin-cleaved aptamers during SELEX by size selection and gel purification.** In this method, we considered 5'FAM-labeled reference markers to indicate where invisible traces of cathepsin-cleaved aptamers migrate through the gel, excise this region, and extract trace cleaved aptamers from the gel.

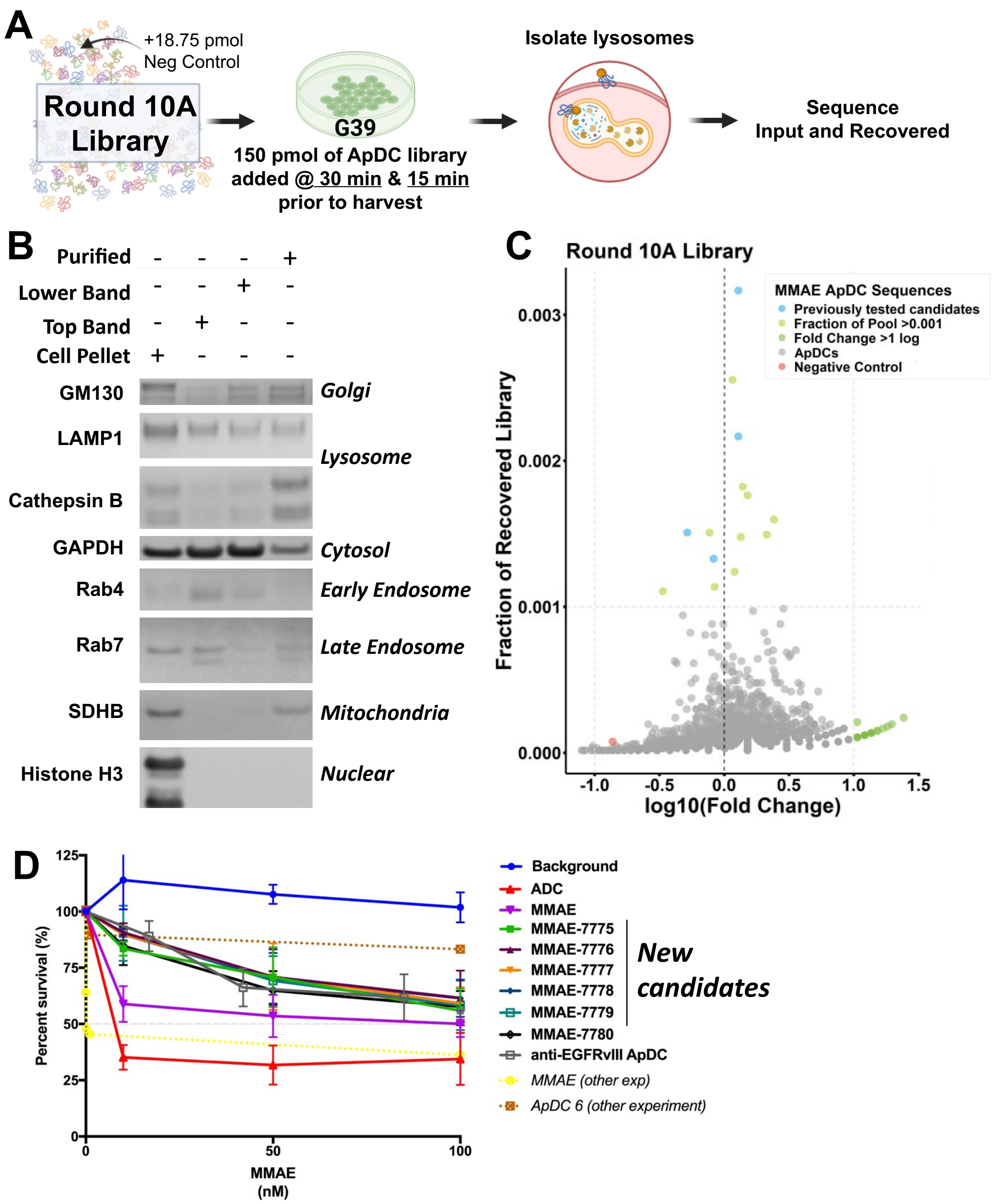


**Supplementary Figure S9. Endolysosomal isolation protocol to identify previous in vivo-selected ApDCs more likely to be internalizing** (7). (A) A schematic diagram of the experiment. (B) Western blot of the extract. (C) Fraction of recovered library vs $log_{10}$(fold change) from input to recovered for individual sequences in the library. (D) Toxicity of the new candidate ApDCs that were selected as they had the largest fold change.

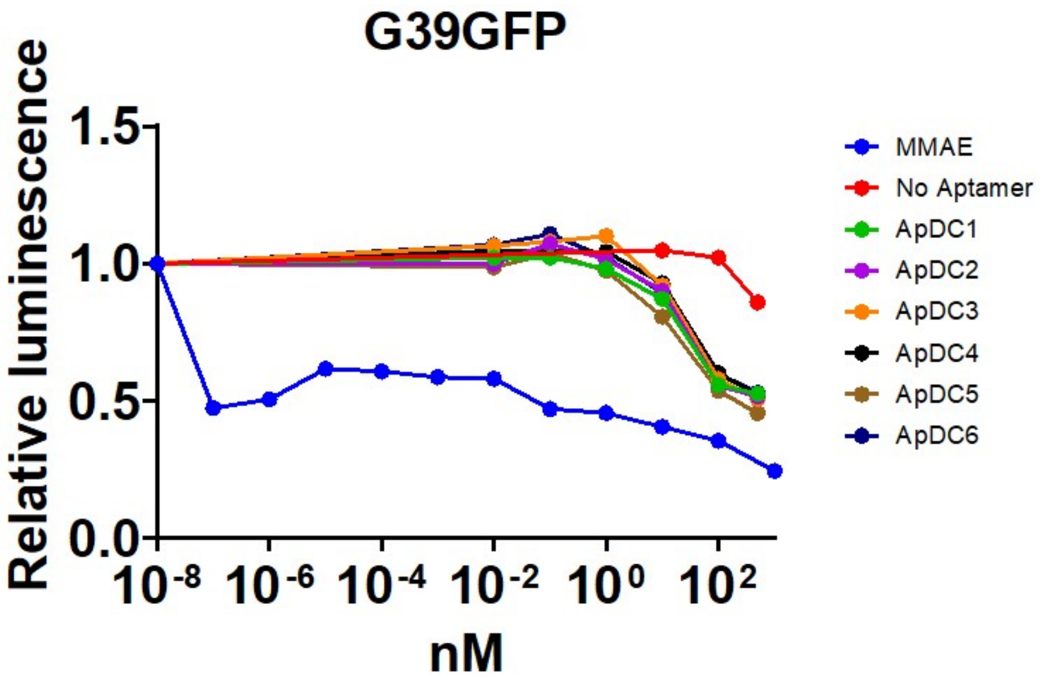


**Supplementary Figure S10.** Lack of toxicity observed previously *in vivo*-selected ApDCs as determined by a CellTiter Glo assay 6 days post-treatment with ApDC, free MMAE or vehicle (7).

A

| First two *in vivo* SELEXs | Third *in vivo* SELEX |
|---|---|
| G39 (BBB very leaky PDX) | G43 (preliminary data suggest **not** leaky BBB) |
| 4 h incubation time for library | 30 or 90 min incubation time for library |
| Reward aptamers in whole “tumor” | Reward aptamers in lysosomal extract from tumor |
| SELEX strategy rewards molecules that accumulate | SELEX strategy rewards molecules that enter a degradation pathway quickly |

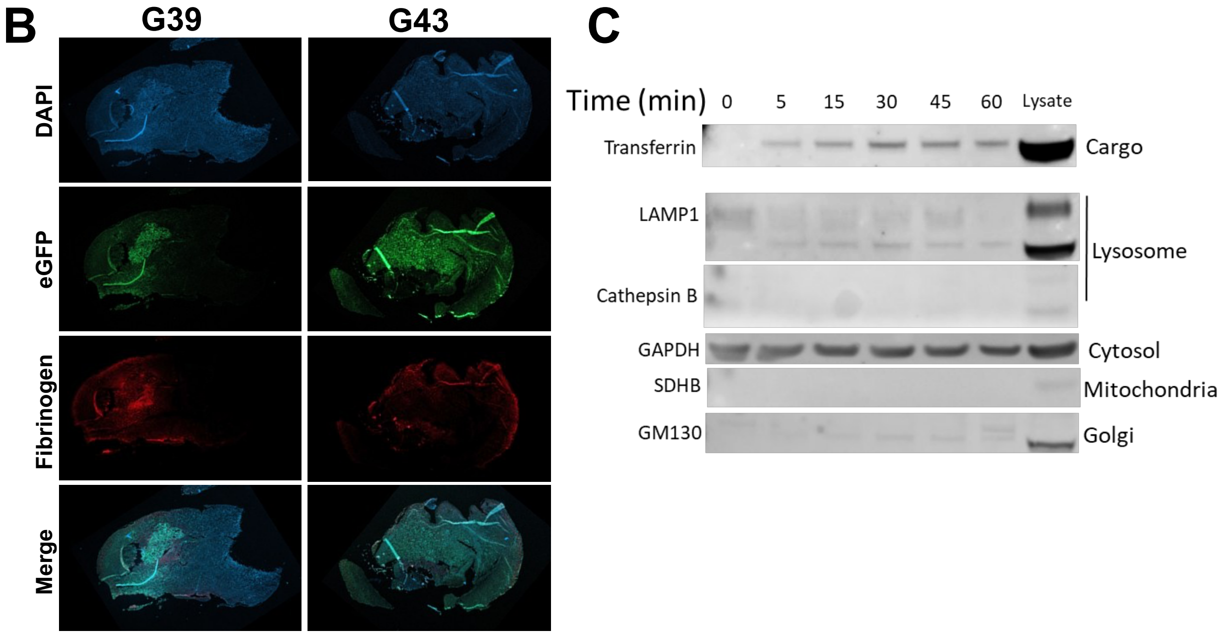


**Supplementary Figure S11. Changes made for this *in vivo* SELEX compared to the first two done by our lab (3, 7).** (A) List of changes. (B) Evaluation of the leakiness of the BBB between G39-eGFP and G43-eGFP. GFP positive staining demonstrates the location of the tumor. Accumulation of fibrinogen in the tumor is a marker of BBB leakiness (2). Note that G43 GFP is on a different scale than G39. The large tumor is evident by DAPI staining, but the GFP signal is much weaker than G39. (C) Fraction of recovered library vs $\log_{10}$(fold change) from input to recovered for individual sequences in the library. (D) Toxicity of the new candidate ApDCs that were selected as they had the largest fold change.

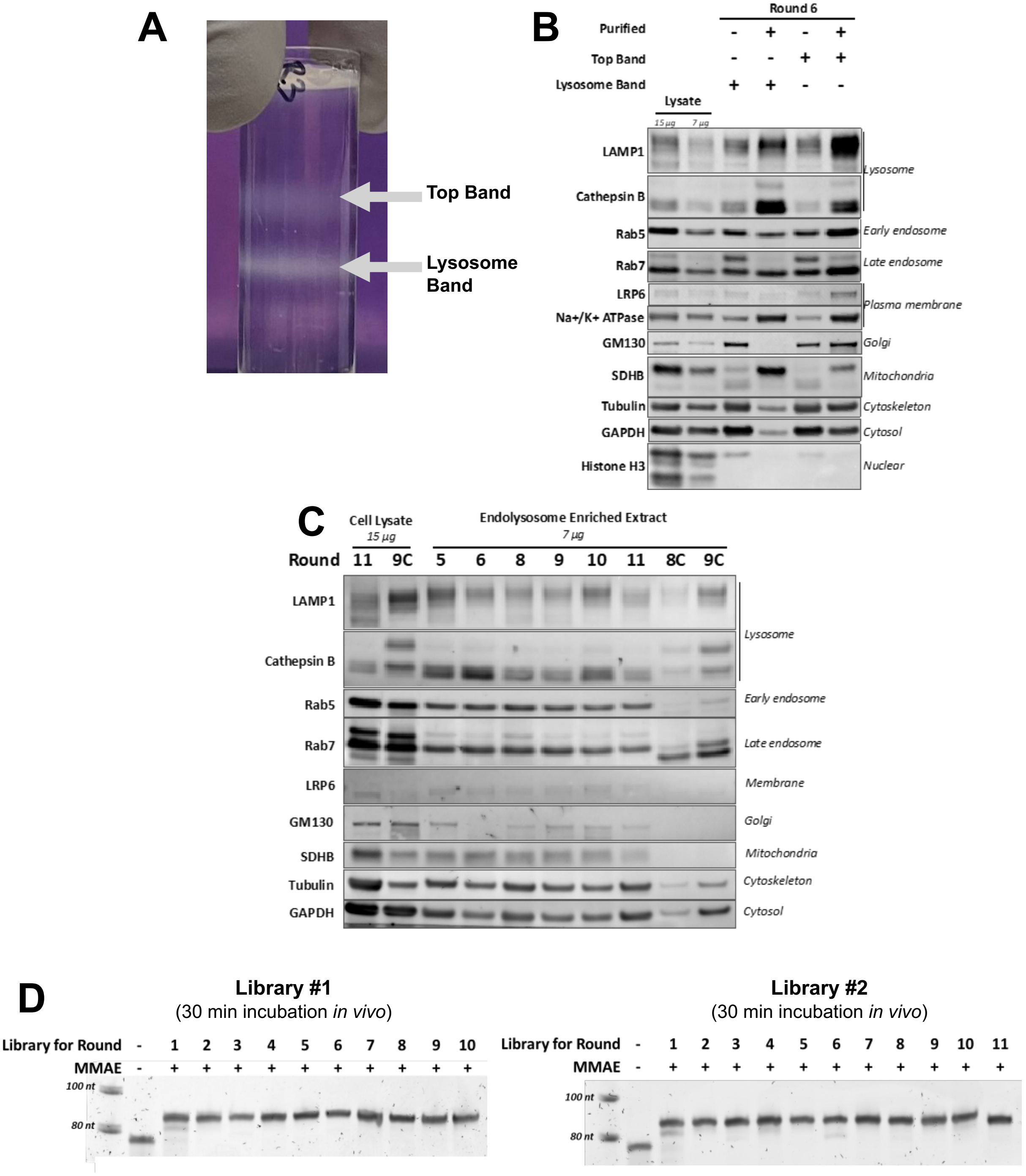


**Supplementary Figure S12. Evaluation of the endolysosome extraction efficiency and library quality during SELEX.** (A) Protein bands after density gradient centrifugation (Abcam Lysosomal Extraction kit) to enrich an endolysosomal fraction. (B) Western blot evaluation of the top and bottom protein bands from (A). (C) Western blot of the recovered endolysosomal fractions during SELEX. "C" denotes round in the case of hybrid selection. (D) Denaturing gel analysis of 80-nt *in vivo* SELEX-only libraries. MMAE conjugation results in mobility retardation comparable to ~10 nt length increase.

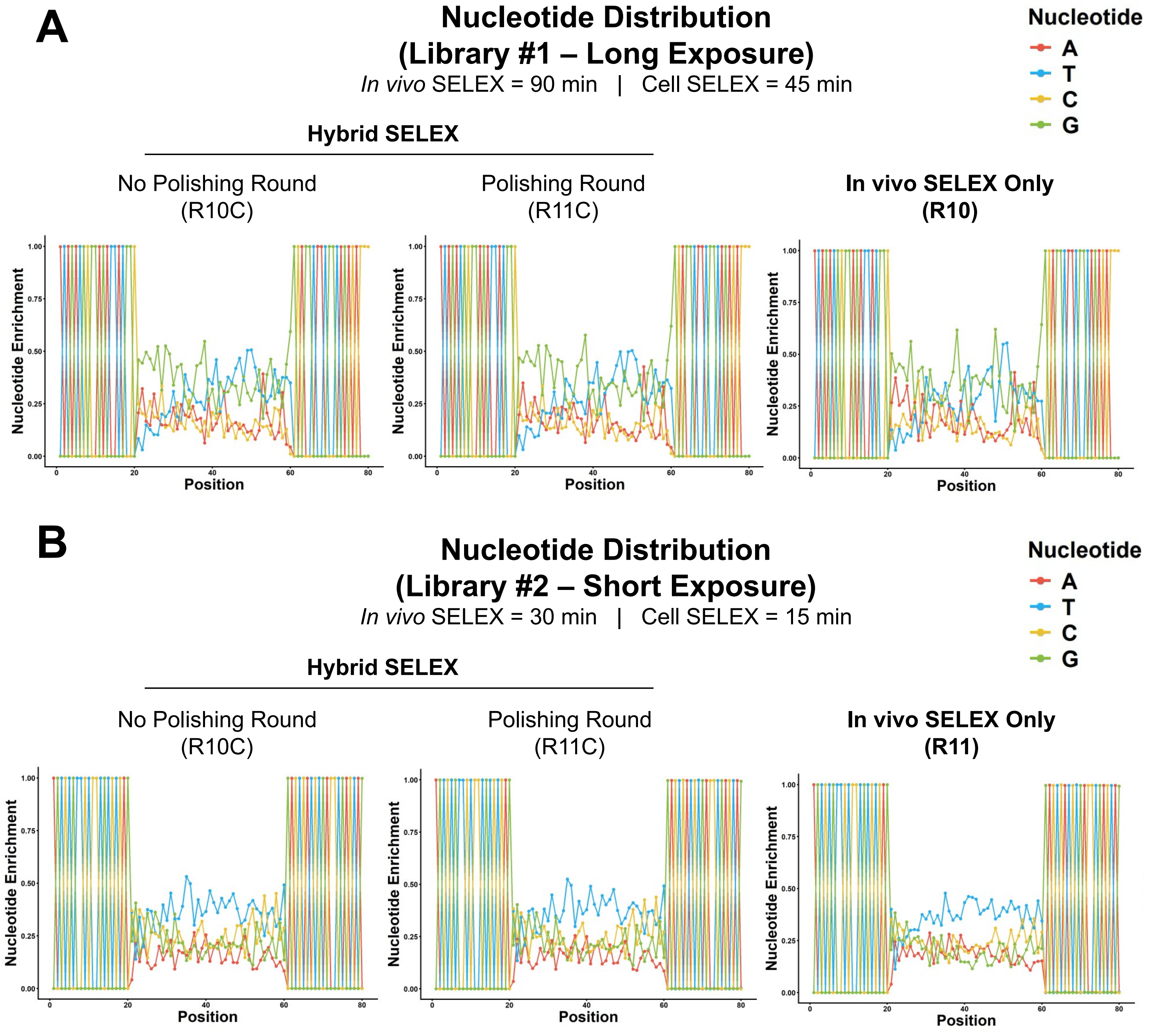


**Supplementary Figure S13.** Final nucleotide distributions in Library #1 (A, long exposure) or Library #2 (B, short exposure).

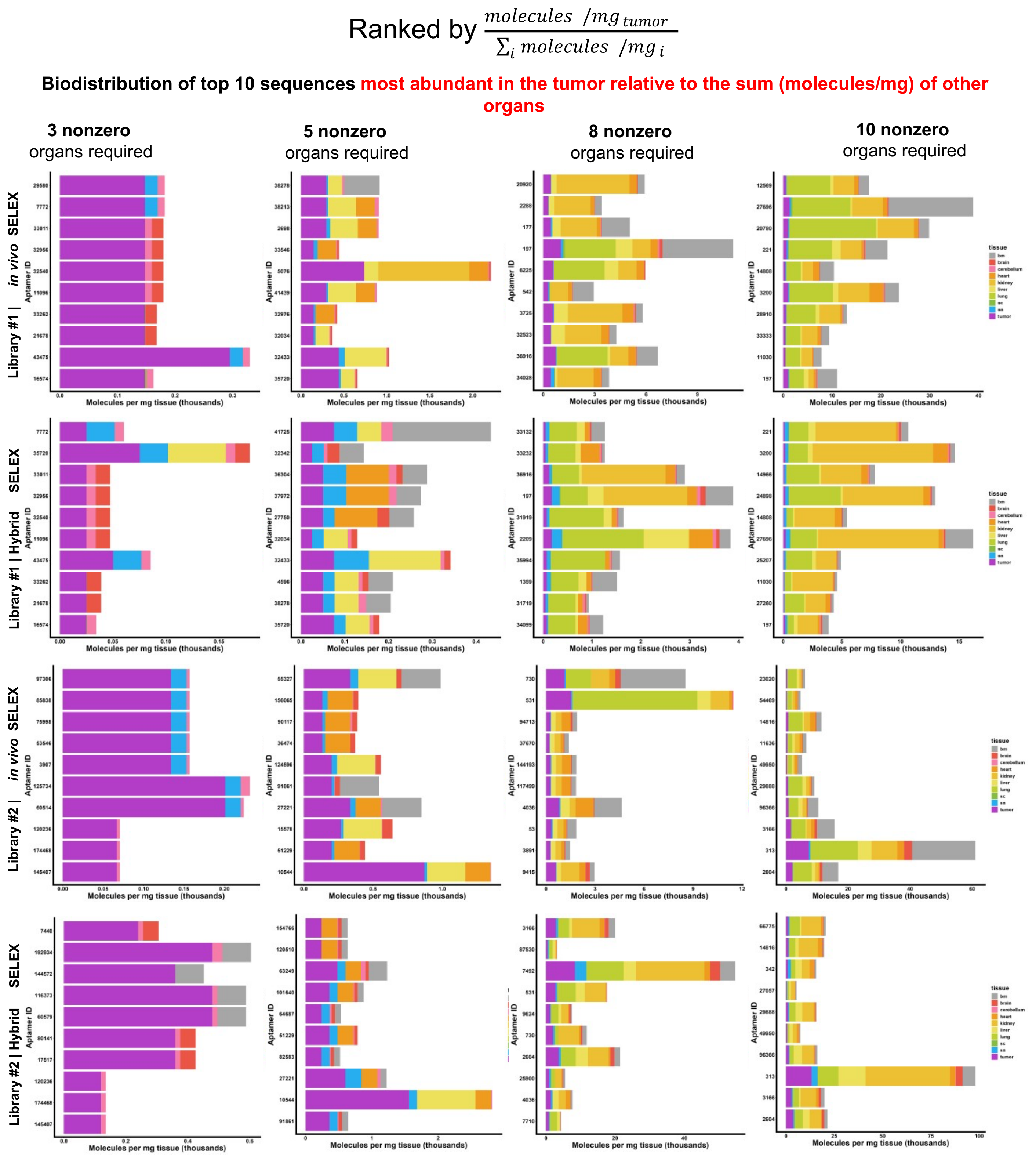


**Supplementary Figure S14.** Attempts to identify rare aptamers in sequencing data with improved tumor homing by ranking molecules by α, where $\alpha = \frac{molecules/mg_{tumor}}{\sum_i molecules/mg_i}$.

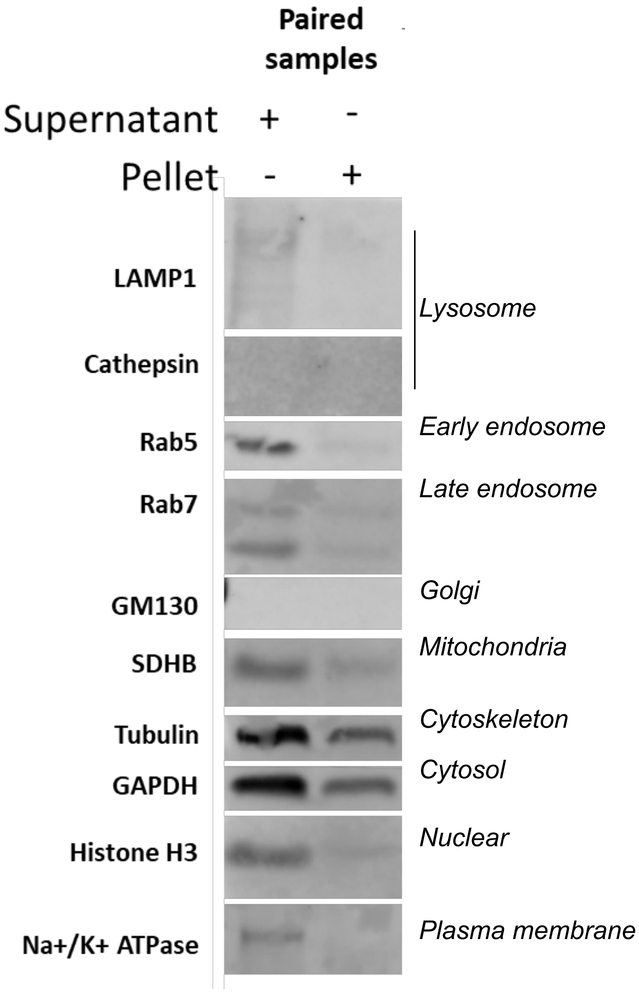


**Supplementary Figure S15. Western blot of sample obtained from bulk tumor processing**. In this protocol, the tissue is homogenized in Buffer P1 with a plastic pestle and then equal volumes of lysis buffer are added to the sample and incubated on ice for 3h. The sample is sonicated three times, heated at 95ºC and subjected to centrifugation at 13.1K × g. The supernatant is analyzed in lane 1 and the pellet resuspended in RIPA buffer, incubated and run in lane 2. Protein concentrations were quantified and normalized to 15 µg per lane. The supernatant is used for sequencing and library preparation in our previous protocol.

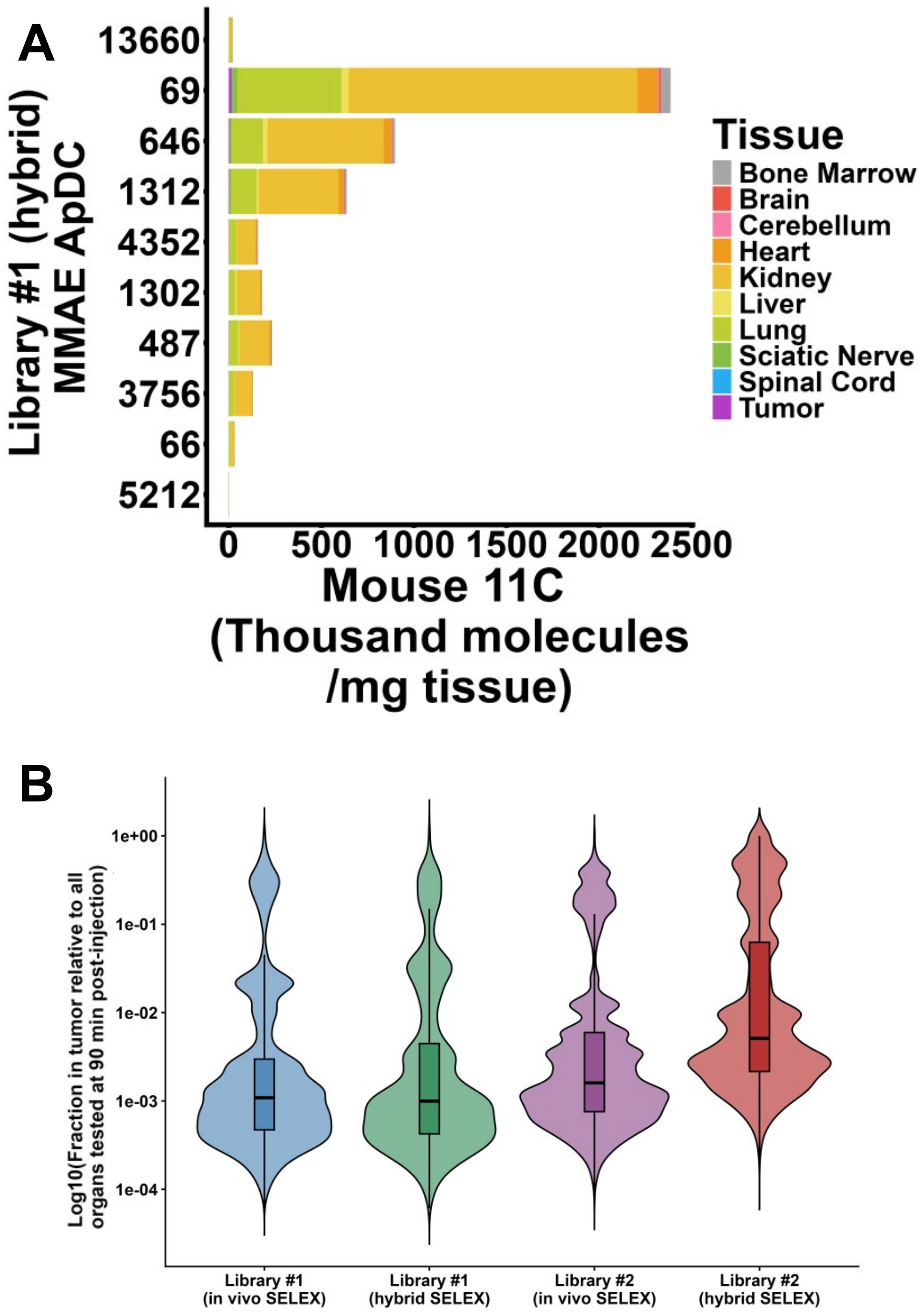


**Supplementary Figure S16. Aptamer biodistributions after hybrid SELEX.** (A) Aptamer biodistributions at the final round by deep sequencing of top sequences in the endolysosomal fraction for Library #1 hybrid SELEX. (B) A comparison across all libraries at the final round for aptamer fraction in tumor relative to all organs.
